# Ohno-miRNAs: intragenic miRNA pairs derived from whole-genome duplication

**DOI:** 10.1101/2025.01.16.632905

**Authors:** Leonardo Agasso, Ivan Molineris, Michele Caselle

## Abstract

Two rounds of whole-genome duplication (WGD) occurred about 500 million years ago and played a major role in the evolution of the vertebrate genomes. Human genes derived from WGD are called “ohnologs”. Ohnologs are involved in fundamental biological processes and significantly contributed to the complexity of the human gene regulatory network. Given the central role of miRNAs in gene regulation, we investigated the contribution of ohnolog miRNAs (Ohno-miRNAs) to the human gene regulatory network. We worked on the identification of miRNA pairs in the human genome derived from the two rounds of whole-genome duplication in early vertebrate lineage and their role in the structure of the human gene regulatory network. By focusing on intragenic miRNAs within ohnolog gene pairs, we identified Ohno-miRNAs as having higher retention rates and sequence similarity compared to miRNAs hosted on paralogue gene pairs from small-scale duplications (SSD). They also show a stronger tendency to regulate common target genes. Analyzing the role of Ohno-miRNAs in the human gene regulatory network, we showed that Ohno-miRNAs are statistically overrepresented in specific network motifs commonly associated with redundancy and complexity, highlighting their central role in this particular layer network.

## Introduction

Gene duplication is the evolutionary process in which a region of DNA hosting a gene produces one or more copies that are temporarily relieved of selective pressure and may potentially develop adaptations for new functions over time. Small-scale duplications (SSDs) include all those processes that duplicate small portions of a genome, usually a single gene or a small set of genes. Since SSDs induce small changes in the genotype, the results on the phenotype are usually of limited magnitude. Whole-genome duplication (WGD), often referred to as “polyploidization”, is instead a process of genome duplication that generates additional copies of the entire genome as a product of nondisjunction during meiosis. WGD events provide raw genetic material to facilitate phenotypic evolution and drastically increase genome complexity. Because of their extreme nature, WGD events are almost exclusively evolutionary dead ends in evolution as they typically involve sudden and dramatic phenotypic changes and are likely to have an immediate impact on the fitness of the organism, compromising both its fertility and short-term survival. Notwithstanding this, whole-genome duplications, although rare, played a major role in evolution. In particular, it is by now accepted the occurrence of two rounds of WGD at the beginning of the vertebrate lineage, as first predicted by Japanese biologist Susumu Ohno in his milestone book Evolution by Gene Duplication [1]. In fact, under particular circumstances WGD events can provide immediate evolutionary benefit to the affected lineage by inducing successful responses to abrupt changes in the environment and, at the same time, they can significantly boost the size and the complexity of the impacted genome causing beneficial effects in the long term [2, 3]. In the human genome, approximately 20-30% of the protein-coding genes can be traced back to WGD events [4]. Thanks to a detailed reconstruction of the evolutionary tree of the vertebrate lineage, a reliable list of putative pairs/quartets of ohnolog genes in vertebrates was recently proposed in [5] and in [6]. A careful analysis of this list allowed us to detect some features which are peculiar to ohnolog genes and may help to shed some light on the mechanism behind ohnolog gene retention. In mammals, ohnologs appear to undergo fewer subsequent small-scale duplication compared to non-ohnolog genes, and seem to be refractory to copy number variation in the human population, compared to genes that have experienced SSD in the vertebrate lineage. These observations suggest that they are likely to be more sensitive to relative quantities (i.e., they are dosage-balanced). Ohnologs are also more frequently associated with human diseases such as the Down syndrome (75% of reported candidate genes for this syndrome are ohnologs) [5]; in addition, ohnolog genes are reported to be three times more likely than non-ohnolog genes to be involved in autosomal dominant diseases and cancer [7]. Ohnolog genes are more likely to be essential (i.e., their removal results in a lethal or sterile phenotype) than SSD-derived paralogues, both as pairs and as singletons [5]. Since these results are at odds with the expected back-up role of duplicated genes (widely observed in less complex eukaryotes) which should provide a buffer against such effects, it is now widely accepted that among them there is a predominance of dosage-balanced genes [5]. From a functional, non-pathological, perspective, ohnologs are found to be more frequently involved in signaling, development and transcriptional regulation [8] and are enriched in Gene Ontology categories associated with the general level of complexity of the organism [9]. From a gene expression point of view, the gene expression profile and subcellular localization display more divergence between the two members of a WGD pair than for members of SSD one [8]. Whole genome duplication has also been shown to affect the structure of the human gene regulatory networks: motif analysis of such networks indicates that the two rounds of WGD have largely contributed to the regulatory redundancy, promoted synergy between different regulatory layers and generated motif that are usually associated with complex functions [10]. One of the goals of the present paper is to improve our understanding of the role played by WGD in shaping the vertebrate regulatory network by including in the game also the miRNA layer of WGD.

Micro-RNAs (miRNAs) are a class of small endogenous, single-stranded, non-coding RNAs (ncRNAs) that play important roles in regulating eukaryotic gene expression, mainly at the post-transcriptional level. In their mature form, miRNAs have a length that can vary between 19 and 25 nucleotides and regulate translation by binding the 3’ untranslated regions (UTR) of the target messenger RNAs (mRNAs), resulting in inhibition of translation or degradation of the mRNA. The miRNA-mRNA interaction is of a combinatorial nature: a single miRNA can directly target hundreds of mRNAs, while a single mRNA can be targeted by multiple miRNAs. MiRNAs typically inhibit the translation of the target transcript by binding to the 3’ UTR, in mammals the binding is strongly dominated by a short subsequence of the mature miRNA, the so-called “seed region” [11], usually consisting of 6–8 nucleotides, mostly situated at positions 2-8, at the 5’ end. More than 60% of human protein-coding genes harbor predicted miRNA target sites [12] and at least 30% of the human protein-coding genome is estimated to be regulated by miRNAs. Aberrant expression of miRNAs is known to play an important role in many pathological processes like allergic diseases [13] and autoimmune diseases [14, 15]The list of WGD genes reported in [5, 6] only contain protein-coding genes. The main goal of the present paper is to (at least partially) extend this list to WGD miRNAs. We shall refer to these miRNAs in the following as Ohno-miRNAs. Their identification opens the way to study the role of WGD in shaping post-transcriptional regulation. We shall see that several of the features already observed in the gene regulatory network at the transcriptional level [10] are present also at the post-transcriptional one and that Ohno-miRNAs are involved in a set of complex network motifs which seem to be precisely associated with their WGD origin.

## 1 Results

### 1.1 Retrieving of putative intragenic Ohno-miRNAs

The primary aim of this paper is to expand upon the list of ohnolog genes identified in [5] and [6] by proposing a catalog of putative intragenic Ohno-miRNAs—pairs of miRNAs that were duplicated during the whole-genome duplication (WGD) events at the origin of the vertebrate lineage in the human genome. To the best of our knowledge, there is currently no comprehensive compendium of ohnolog non-coding genes in the literature.

To achieve this, we build on existing knowledge of WGD protein-coding genes [5, 6], searching for pairs of duplicated intragenic miRNAs located within pairs of ohnolog protein-coding genes. This approach enables us to analyze approximately 50% of the known miRNAs in the human genome, as nearly half of them are recognized as intergenic (see Supplementary Material, Section 2). This observation provides strong evidence that intragenic miRNAs were likely duplicated during the same WGD events as their host genes. Another possibility that could result in such a configuration involves the following sequence of events: after the WGD, two ohnolog copies of an ancestral protein-coding gene G (named G1 and G2) are retained; subsequently, (1) a new miRNA emerges within G1, (2) this miRNA undergoes a small-scale duplication (SSD), and (3) the duplicated copy is inserted precisely into G2. This scenario is significantly more complex and less probable than the simpler hypothesis of miRNA duplication during WGD, and thus we chose to disregard it. We retrieved 8,089 protein-coding genes forming 10,271 WGD-derived pairs (ohnolog pairs) from existing databases of ohnolog genes [5, 6]. From the Ensembl database we downloaded 13,784 protein-coding genes forming 122,863 paralogue pairs that are not ohnologs and that we shall define in the following as SSD pairs (see the methods section for a detailed discussion of these selections). In terms of single genes, there’s a large intersection between the two sets, in fact 6,048 genes are involved both in WGD and SSD duplication.

We then downloaded from GENCODE all the genes annotated as miRNA and looked for all the instances in which *both* genes of a WGD or SSD pair host a miRNA whose strand orientation is conserved. We selected all the intragenic miRNAs, including the intronic ones and those located in the exons (see fig.2). When more than one miRNA was present in one of the two ohnolog genes we leveraged sequence similarity to select just one pair (see methods). Only a fraction of the human miRBase entries are strongly supported miRNA genes [16] and false positives are most probably propagated in datasets such as GENCODE [17] whose miRNA annotations use miRBase data. To address this problem we selected for our subsequent analysis only 505 miRNA genes for which at least one primary transcript is recognized as *bona fide* by MirGeneDB [18], a curated database of miRNA genes that have been manually validated and annotated. We report for completeness the analysis performed on the whole set of miRNAs in the Supplementary Material, the results we obtain in this case are in line with what we observe in the main analysis.

We identified 20 pairs of miRNAs hosted on ohnolog genes and 33 pairs hosted on SSD pairs; moreover we classified MIR196A1-MIR196A2 as Ohno-miRNAs after manual curation (see Methods). These pairs are listed in the Supplementary Material (Table 1 and Table 2). As a last step we selected only those pairs of miRNAs recognized by Ensembl as paralogue miRNAs. Remarkably enough, 17 out of the 20 ohnologs pairs survived this selection, while on the SSD side only one out of the 33 pairs survived. We list in tab. 1 and tab. 2 our results. We consider the miRNA pairs listed in tab.1 as a highly reliable list of Ohno-miRNA pairs.

**Table 1.**
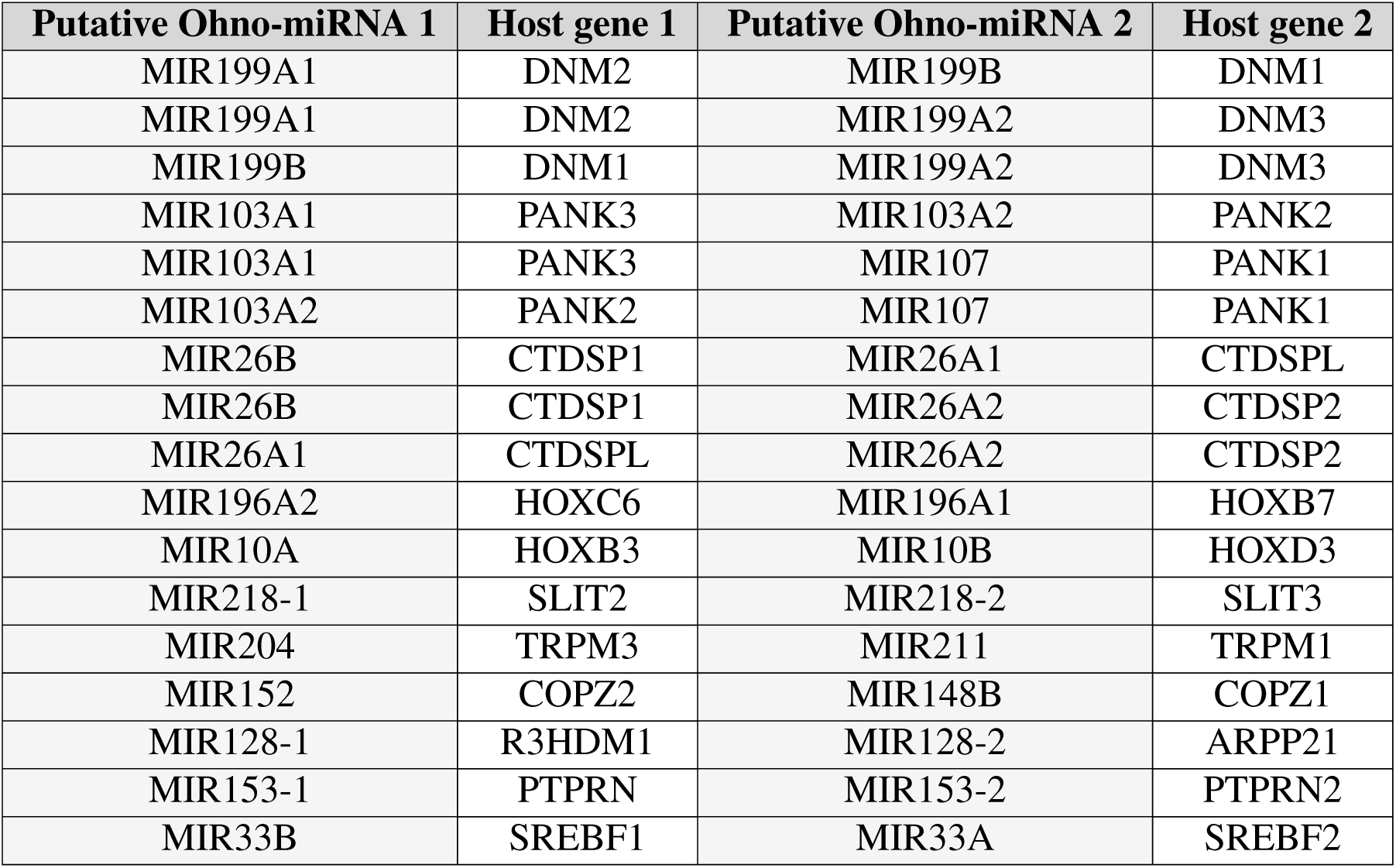
Putative intragenic ohnolog miRNA pairs with their host genes.

**Table 2.**
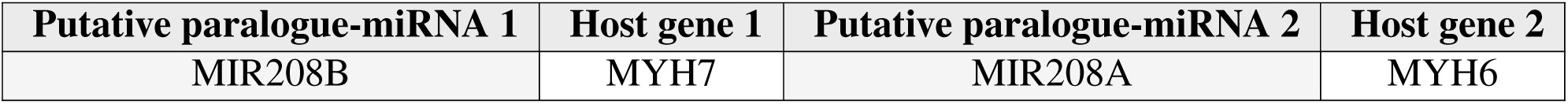
The only pair of putative intragenic SSD-derived paralogue miRNAs surviving our filters, with their respective host genes.

The only remaining instance of a SSD pair of intronic miRNAs is an example of interplay between small-scale and whole-genome duplication (the same two miRNAs: MIR208A and MIR208B are also involved in two WGD pairs with MIR499A, although not recognized as paralogues by Ensembl) which we shall discuss in detail in Section 2.5 below.

Interestingly we observed that many Ohno-miRNA are hosted on gene families, including:

- pantothenate kinase (PANK) genes: MIR103A1 (PANK3), MIR103A2 (PANK2), MIR107 (PANK1);
- C-terminal domain small phosphatase (CTDSP) genes: MIR26A1 (CTDSPL), MIR26A2 (CTDSP2), MIR26B (CTDSP1);
- dynamin (DNM) genes: MIR199A1 (DNM2), MIR199A2 (DNM3), MIR199B (DNM1);
- myosin heavy chain (MYH) genes: MIR208A (MYH6), MIR208B (MYH7), MIR49o9A (MYH7B);
- homeobox (HOX) genes: MIR196A2 (HOXC6), MIR196A1 (HOXB7), MIR10A (HOXB3), MIR10B (HOXD3).

### 1.2 Putative ohnolog miRNAs are more conserved than paralogue ones

Before applying the Ensembl filter, we identify 20 putative intragenic ohnolog miRNA pairs and 33 putative intragenic paralogue miRNA pairs. These pairs are determined based solely on their presence within ohnolog or paralogue protein-coding genes and strand orientation. However, the Ensembl filter, which restricts the analysis to miRNA pairs recognized as homologous by Ensembl, reduces the number of putative paralogues from 33 to just 1, while 17 out of the 20 ohnolog pairs are retained. To better understand the impact of this filter, we analyze the sequences of both sets of miRNA pairs prior to applying it. Putative intragenic ohnolog miRNA pairs display a higher sequence similarity compared to putative intragenic paralogue miRNA pairs, except for the aforementioned MIR208A-MIR208B pair (figure 3-A). A similar pattern emerges when similarity is assessed not through sequence alignment scores, but by evaluating shared target genes (3-B). Many of these putative paralogues are not recognized as homologous by Ensembl, causing a drop in the number of putative paralogues due to the Ensembl filter step of our pipeline (fig.1). This suggests two possible scenarios: (1) the excluded miRNA pairs are actually not paralogues, thus were not duplicated along with their host genes but emerged independently afterward, or (2) these miRNA pairs were duplicated during SSD events but subsequently underwent rapid evolutionary divergence (neofunctionalization), leading to poor sequence similarity and making them difficult to be recognized as paralogues by sequence-based algorithms. The latter scenario is more plausible and highlights a clear difference in the evolutionary trajectories of WGD-derived and SSD-derived pairs. The magnitude of this effect is marked and emphasizes the importance of miRNA regulation in the vertebrate lineage and the evolutionary pressures shaping these distinct classes of miRNAs.

**Figure 1.**
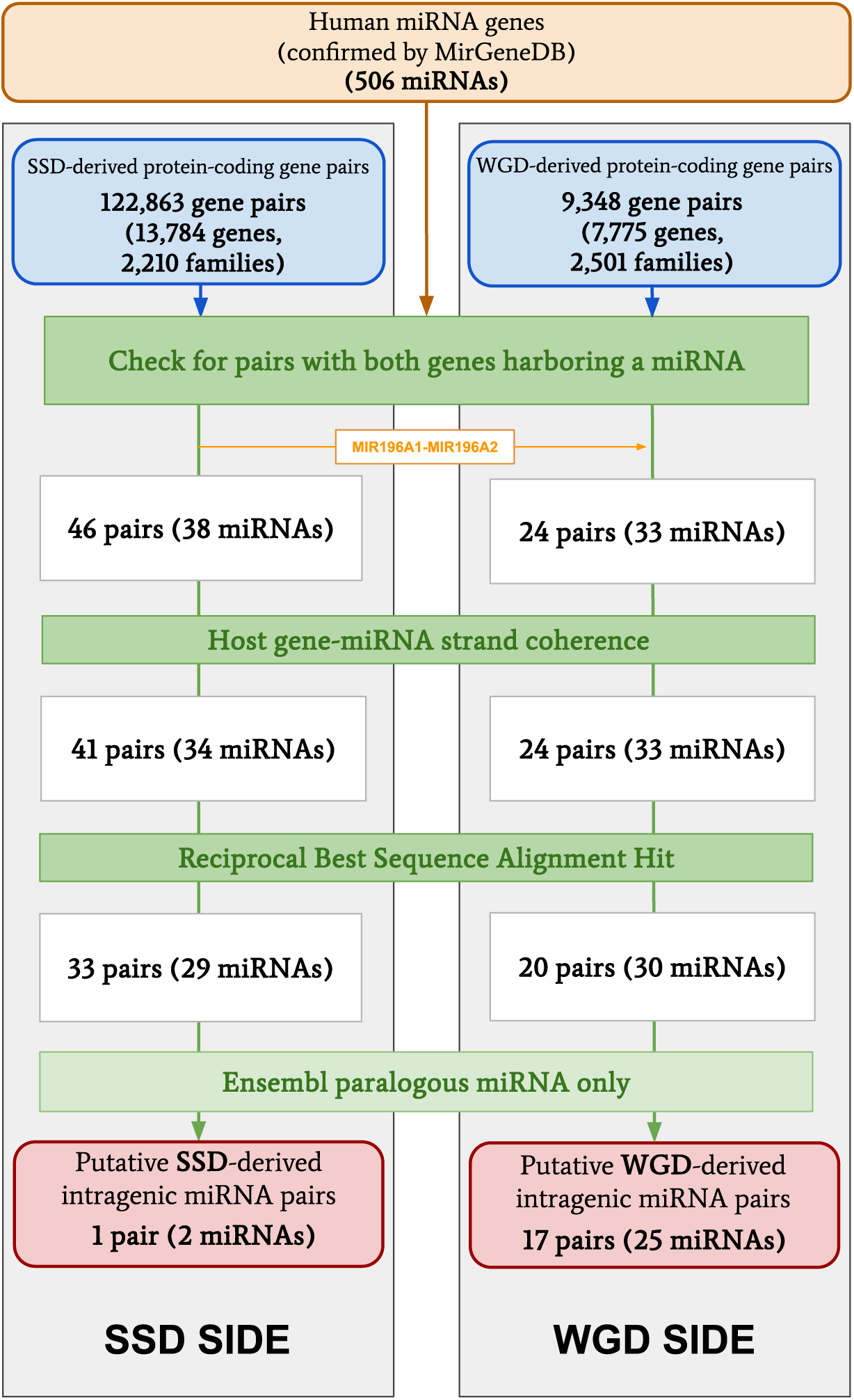
Scheme of the pipeline followed to retrieve putative intragenic Ohno-miRNA pairs.

**Figure 2.**
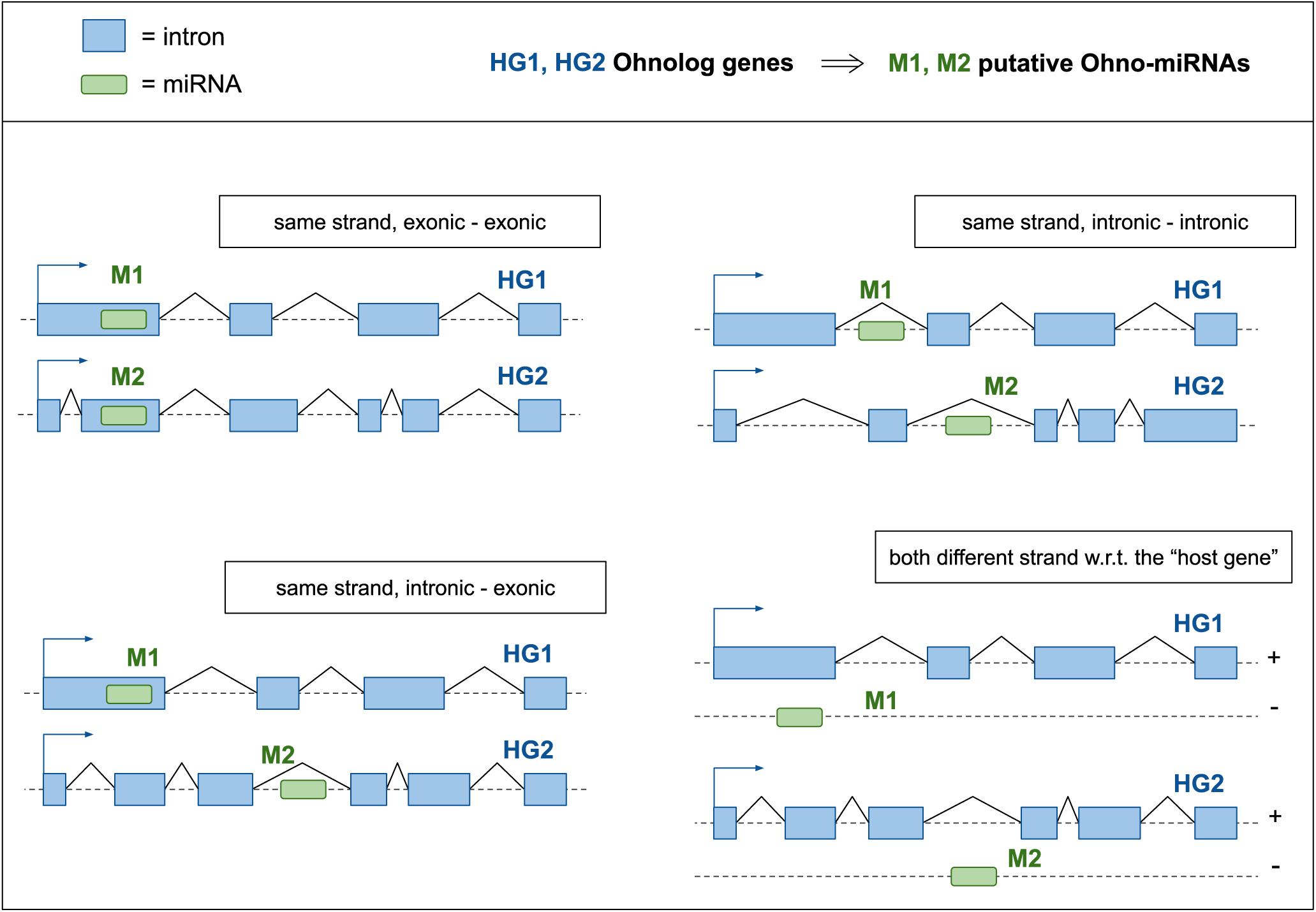
Schematic representations of all the different configurations of putative Ohno-miRNAs in relation to ohnolog gene pairs.

**Figure 3.**
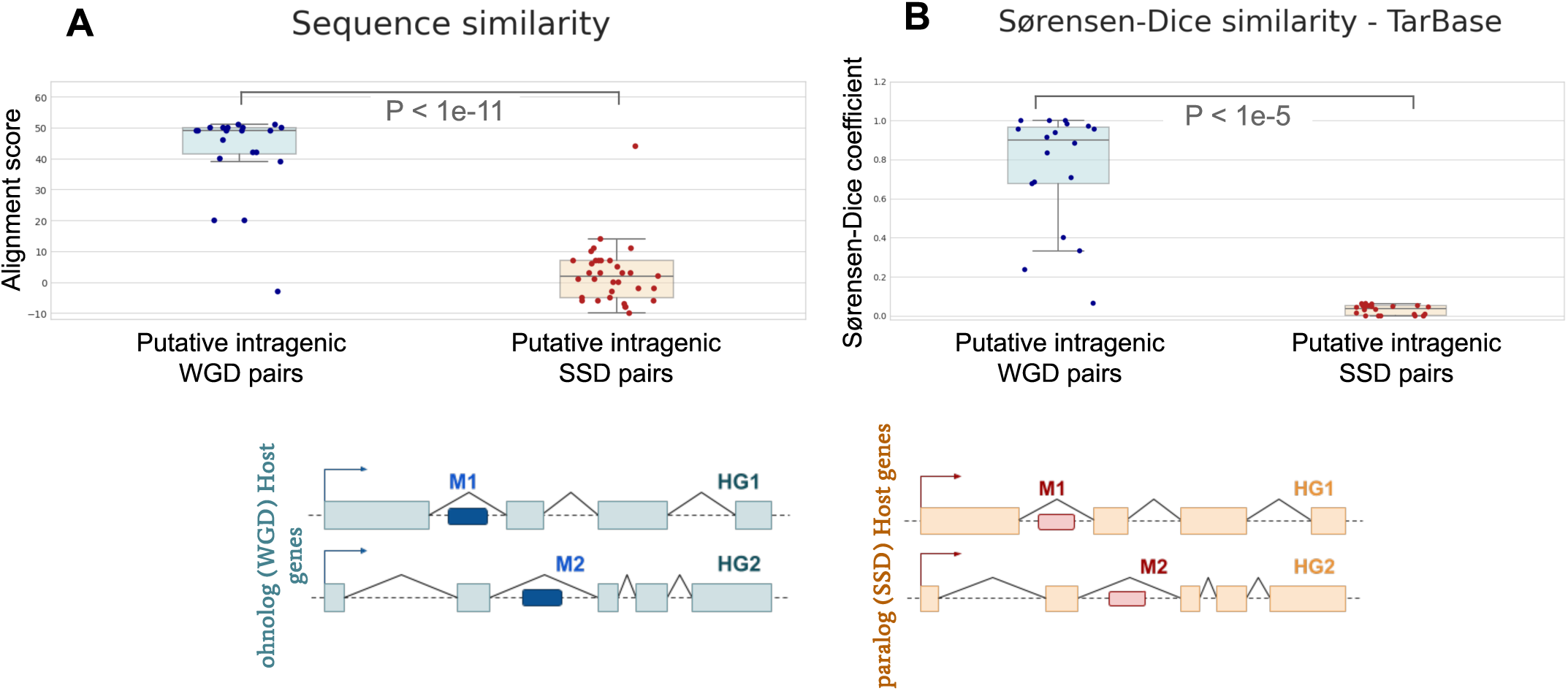
Comparison of alignment scores (miRNA sequence similarity) for WGD and SSD pairs (left panel) and Sørensen-Dice coefficients (target similarity) in the TarBase network (right panel) before applying the Ensembl filter. Both metrics show statistically significant differences between WGD and SSD pairs (Kolmogorov-Smirnov test). Some pairs are absent from panel B due to the lack of experimental data in TarBase.

#### 1.2.1 Conservation of putative ohnolog miRNAs is confirmed in the mouse genome

The conservation of putative ohnolog miRNAs can be checked across different vertebrate species. To validate this conservation, we analyzed the mouse (*Mus Musculus*) genome using the same pipeline described in Fig. 1. The results further support the trends observed in the human genome, demonstrating that intragenic ohnolog miRNAs retain higher sequence similarity compared to their SSD-derived counterparts. Fig. 4 shows the results of the sequence alignment in the mouse genome.

**Figure 4.**
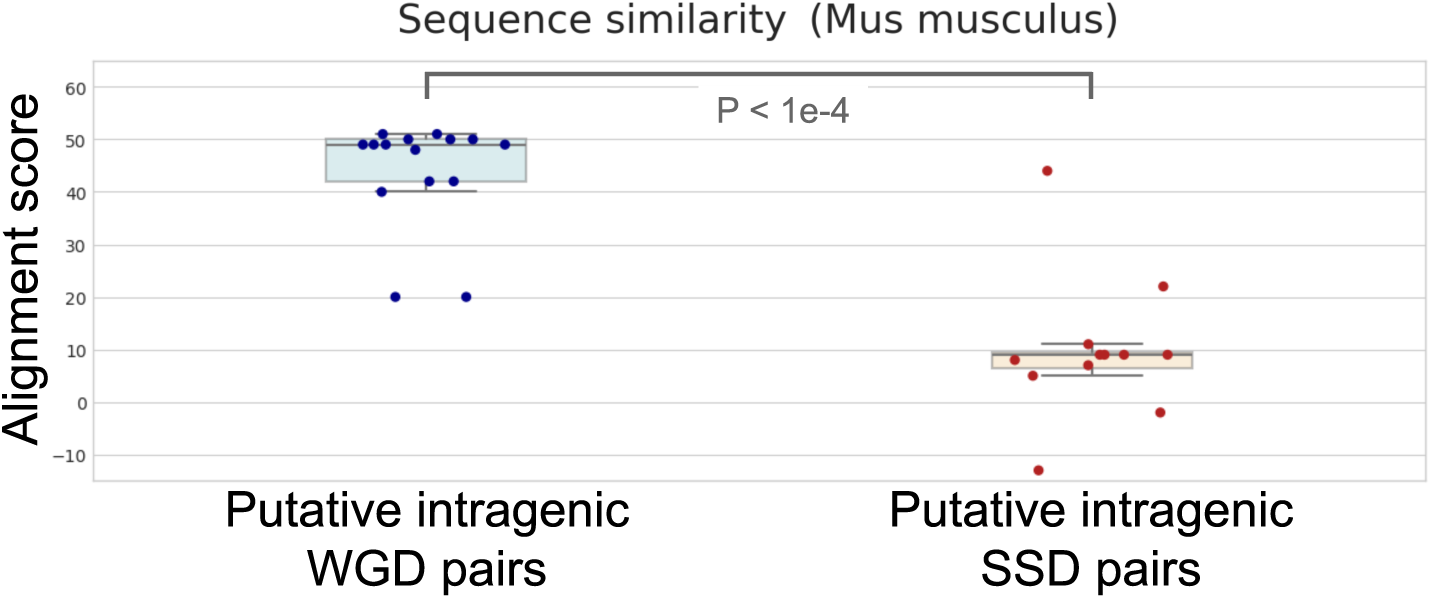
Comparison of alignment scores (miRNA sequence similarity) for WGD and SSD pairs for intragenic miRNA pairs detected in the mouse genome following the pipeline described in Fig.1. The distributions of alignment scores for both WGD and SSD pairs exhibit statistically significant differences (Kolmogorov-Smirnov test), consistent with the patterns observed in the human genome.

We report and discuss the list of putative intragenic Ohno-miRNAs in the mouse genome in the Supplementary Material, Section 7.

### 1.3 Ohno-miRNAs show peculiar patterns within the regulatory network

Equipped with the list of Ohno-miRNAs reported in tab.1 we can now study the role that they play in the human regulatory network. This study is the natural continuation of the analysis of ref. [10], where the role of WGD transcription factors in shaping the human regulatory network was highlighted. In [10] it was shown that ohnolog pairs of transcription factors were involved in a few specific regulatory motifs which were detected looking to their enrichment in the regulatory network with respect to suitable null models. The aim of this section is to extend this analysis to miRNAs. To address this challenge we used two different miRNA-target networks, TarBase [19] and MiRDIP [20], and two different protein-protein interaction networks, PrePPI [21] and STRING [22]. We report in the main text only the results obtained with the TarBase and PrePPI databases and in the Supplementary Material (Sections 3,4), those obtained with the MiRDIP and STRING ones. We find essentially the same results even if the dataset used as references is biased in opposite directions and with limited overlap (see Discussion).

#### 1.3.1 The out-degree distribution of Ohno-miRNAs slightly differs from that of ordinary miRNAs

As a first step, we analyzed the out-degree distribution of the 28 putative Ohno-miRNAs present in the TarBase network, comparing it with that of SSD-duplicated miRNA and all the remaining miRNAs in the network. In this context, the out-degree represents the number of target genes of a given miRNA. We observe that duplicated miRNAs tend to have a higher out-degree than non-duplicated ones, with this tendency being slightly more pronounced in Ohno-miRNAs than in paralogue ones (see fig.5).In all the subsequent analysis we take into account this difference, ensuring that our results are not just due to the observed difference in out-degree distribution. Indeed, all the results reported below were obtained by comparing retrieved networks with a null model generated by randomly reshuffling the links while keeping the degrees of the nodes constant (see Methods). Moreover, this approach accounts for biases such as the TarBase tendency to report more targets for genes that have received greater attention from the scientific community.

**Figure 5.**
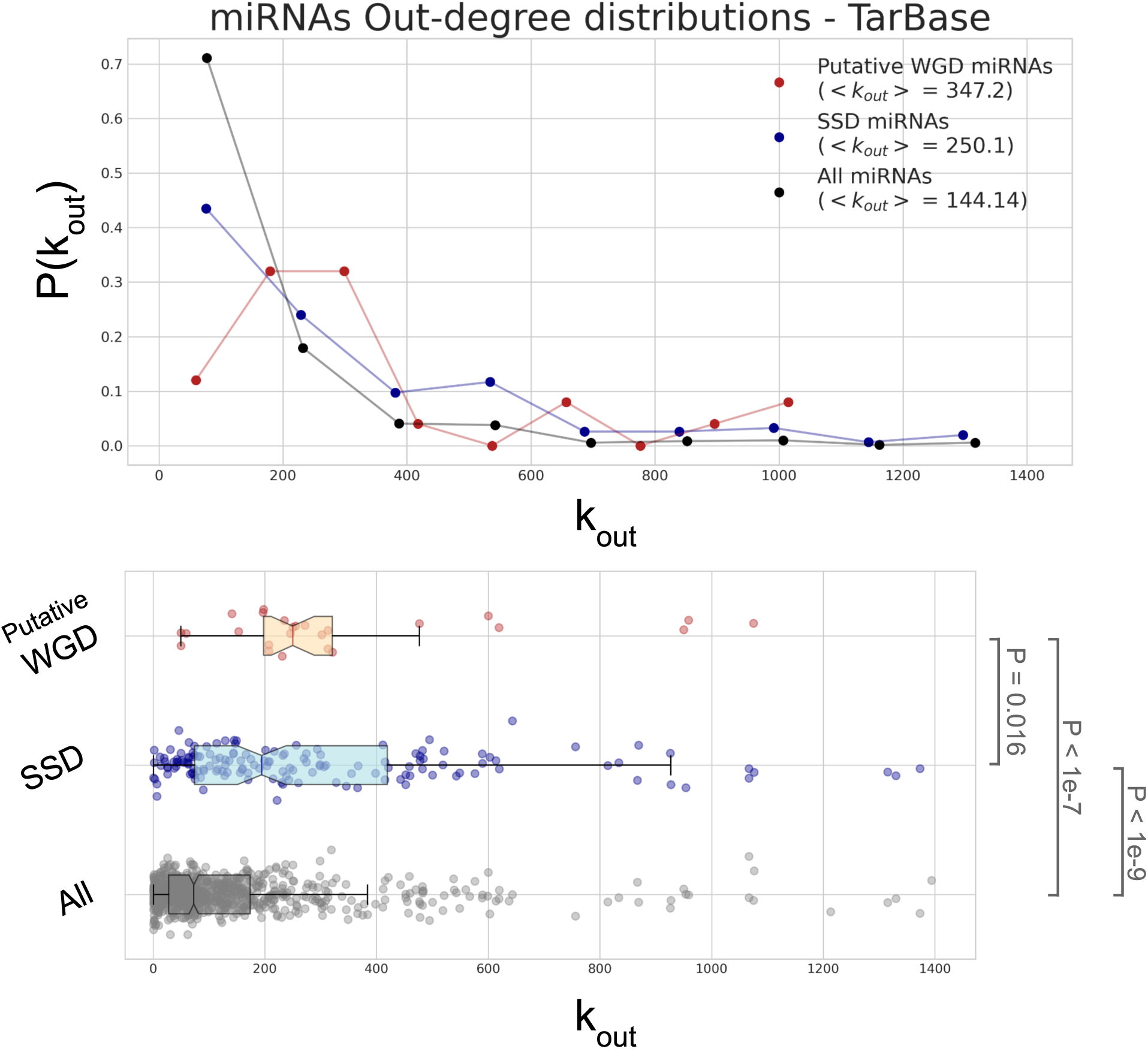
Out-degree distribution of putative Ohno-miRNAs compared with that of the paralogue miRNAs from Ensembl, and all the miRNAs in the TarBase network.

#### 1.3.2 V-motif enrichment in the miRNA-target networks

The first non-trivial motif we analyzed is the so-called “V-motif” consisting of a duplicated pair of miRNA interacting with a common target gene. Ohno-miRNA pairs display a significant tendency to be involved in V-motifs. A general motif enrichment analysis (see Methods, fig.12,A) indicates that both the putative WGD and SSD pairs are enriched in V-motifs, with very high Z-scores: *Z_Putative WGD_ >* 100, *Z_SSD_ >* 300. Such an analysis is not very informative since it does not make it clear whether this enrichment is due to all the duplicated pairs in a set, or just to a subset. To overcome this problem, we evaluated a “pairwise” Z-score (see Methods, fig.12,B).

Observing fig.6, both SSD and WGD pairs exhibit a significant tendency to retain common targets after duplication, thus generating V-motifs. However, we see a clear enrichment of V-motifs in the putative Ohno-miRNA set with respect to one with SSD duplicated ones, which is similar to what was observed in transcription factors [10]. This signal of “redundancy” in gene regulation seems to be a hallmark of WGD and is most likely related to the increase in complexity of the regulatory network of vertebrates compared to other organisms [23]. This observation holds even when removing duplicated miRNA pairs younger than the *Sarcopterygii* clade (the “Ancient SSD” set), that interestingly show low Z-scores.

**Figure 6.**
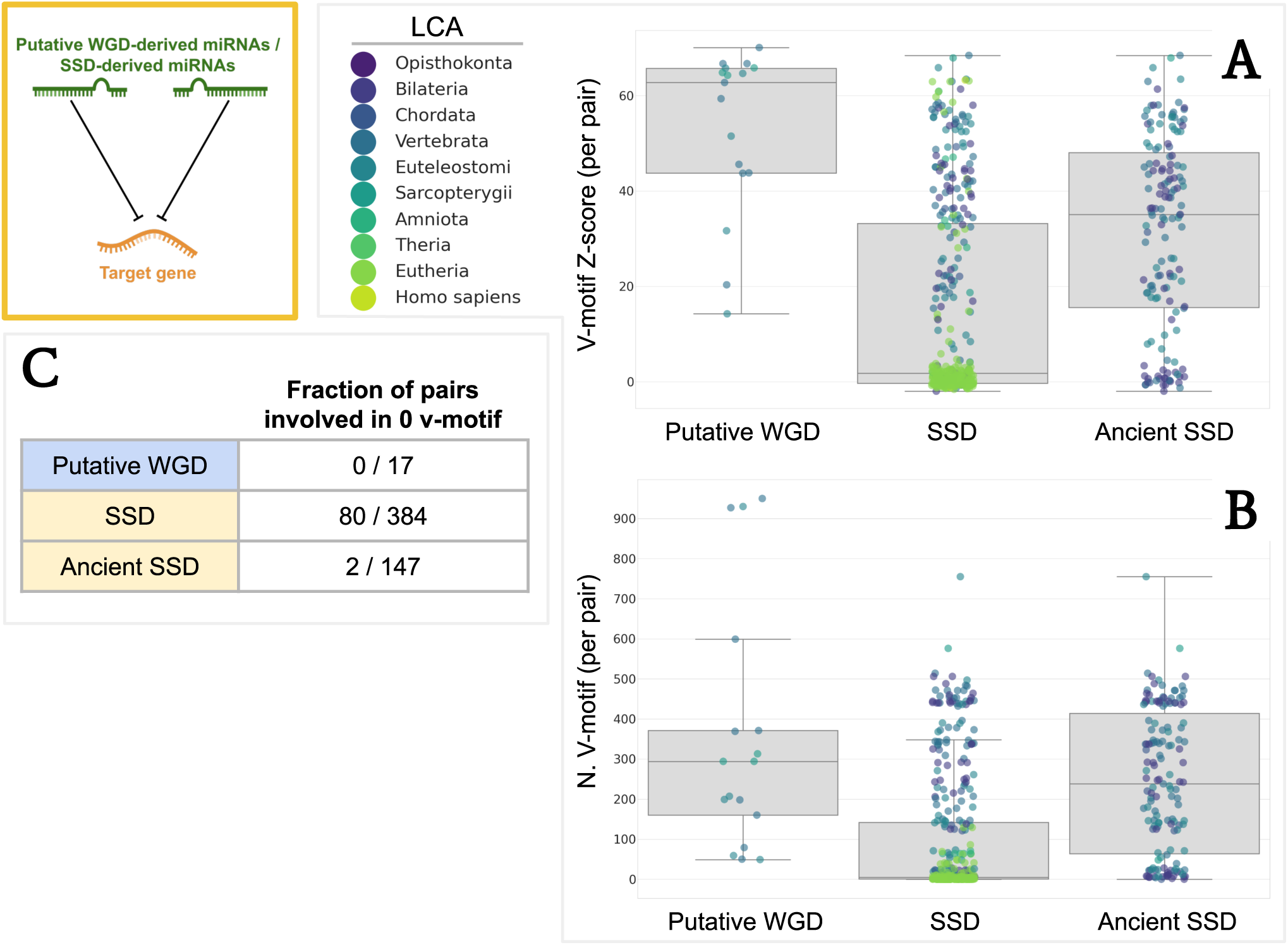
V-motif enrichment in the TarBase network: **(A)** Distribution of the Z-scores refers to the enrichment of each pair with respect to the null model (each dot is a pair, with a color corresponding to the last common ancestor of the pair). The first set (Putative WGD) reports the Z-score of motif enrichment of putative Ohno-miRNA pairs, the second set (SSD) reports the same quantity for all the other pairs of paralogue miRNAs in Ensembl, the third sets (Ancient SSD) reports only SSD pairs whose last common ancestor is older than *Sarcopterygii*. **(B)** Number of V-motif involving each pair. **(C)** Number of pairs not involved in any V-motif for each of the three sets. P-values from the comparison of the Z-score distributions: *P_Putative WGD vs SSD_ <* 1*e−* 7, *P_Putative WGD vs Ancient SSD_ <* 1*e−* 4 (Kolmogorov-Smirnov test).

#### 1.3.3 Bifan enrichment

The second regulatory motif we analyzed is the bifan, a structure in which a pair of paralogue miRNAs simultaneously regulate a pair of paralogue target genes (see for instance [24] for a discussion of the role and functions of this motif at the transcriptional level)^1^.

We evaluated the enrichment of bifans with respect of null models for three sets of miRNA pairs: the “Putative WGD” set (putative Ohno-miRNA pairs), the “SSD” set (paralogue miRNAs identified in Ensembl), and the “Ancient SSD” set (SSD pairs whose last common ancestor is older than Sarcopterygii). As for the V-motifs, both the putative WGD pairs and the SSD pairs are globally enriched strongly in bifans, with *Z_Putative WGD_*= 34 and *Z_SSD_ >* 100 (These very strong enrichment are consistent across different networks as shown in the Supplementary). For the same reasons outlined in the previous paragraph, we switched to a pairwise enrichment analysis.

The results are shown in Fig. 7 In the first set, both the miRNA pairs and their target gene pairs are required to have originated from whole-genome duplications (WGD). Conversely, in the “SSD” and “Ancient SSD” sets, only SSD-duplicated genes and miRNAs are considered. This distinction allows us to isolate bifans likely formed during the WGD event and subsequently conserved.

**Figure 7.**
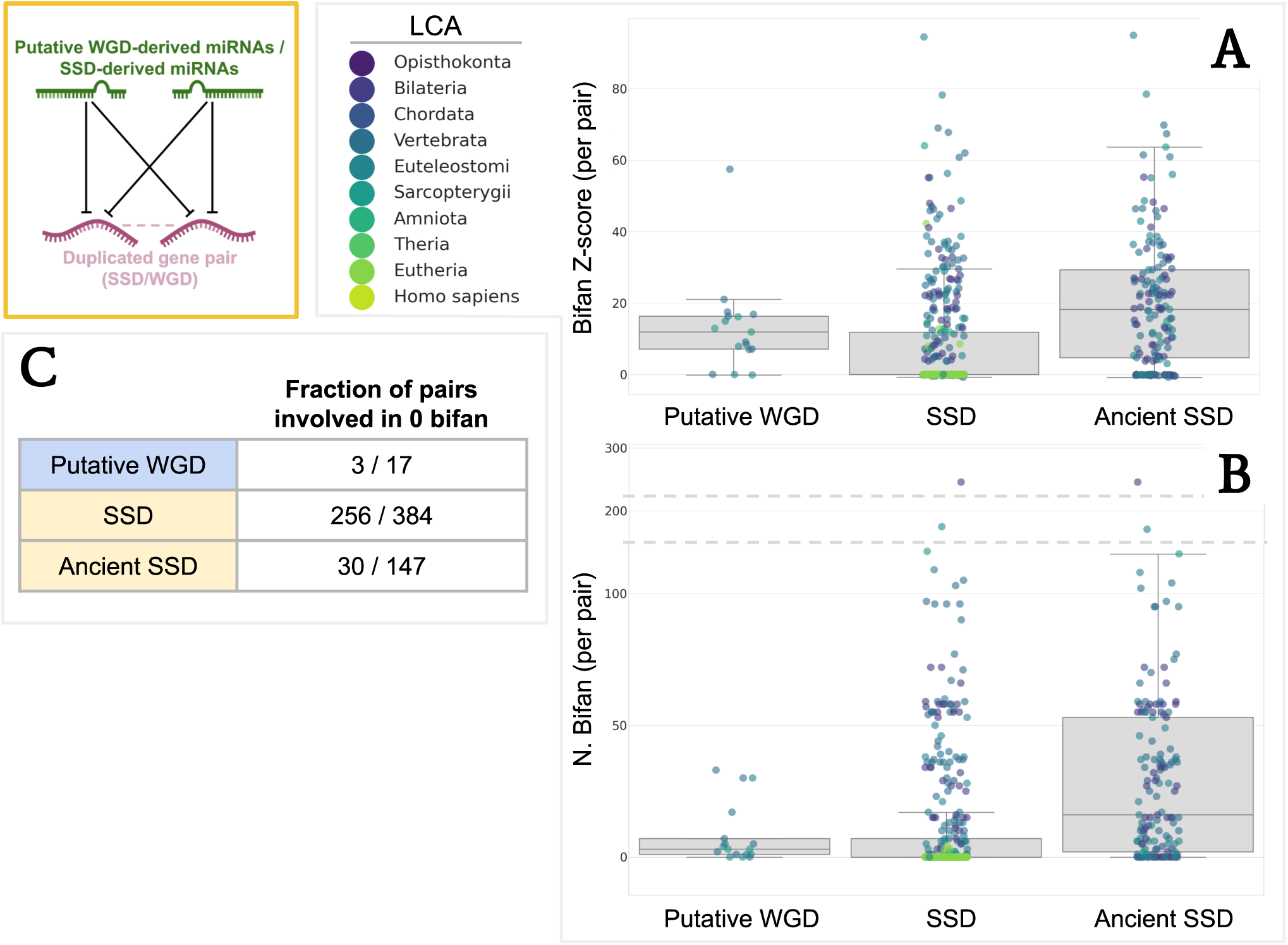
Bifan enrichment in the TarBase network: **(A)** Distribution of the Z-scores refers to the enrichment of each pair with respect to the null model, the definition of the three sets and the color-coding is the same as in Fig.6. **(B)** Number of bifans involving each pair. **(C)** Number of pairs not involved in any bifan for each of the three sets. P-values from the comparison of the Z-score distributions: *P_Putative WGD vs SSD_ <* 1*e−* 4, *P_Putative WGD vs Ancient SSD_* = 0.54 (Kolmogorov-Smirnov test).

Fig. 7,B contains a lot of information. Let us discuss it in detail:

- Observing the Z-score of all the pairs (both of the WGD or of the SSD type) they show a strong tendency to target a pair of duplicated targets (i.e., to create a bifan). Such large Z-scores (fig.7,A) suggest a strong evolutionary pressure to generate these motifs (or to keep them if they are generated during the duplication event). Considering the pairs independently, the distributions of the Z-scores do not differ significantly between the WGD and the SSD-derived pairs.
- At the same time, we see that a lot of SSD-derived miRNA pairs are involved in exactly zero bifans (fig.7,C). This is the main difference between WGD and SSD pairs and is the reason why we see a slight increase in the Z-score distribution of WGD bifans with respect to the SSD ones.
- In the SSD set we see a precise evolutionary pattern: newly duplicated miRNAs typically are not involved in bifans (enrichment of green points at the bottom of the distribution). The simplest explanation of this suggests a non-trivial behavior: the creation of bifans by SSD requires long evolutionary times and is, most likely, a two-step process. First a miRNA is duplicated, the two miRNAs at the beginning have all their targets in common and (as we have seen above) there is a strong evolutionary pressure to keep this redundancy of targets. Second, one of these common targets is duplicated, thus generating a bifan which is then conserved. This explains the difference between The WGD and SSD sets: during the WGD event *both* the miRNA and the target are simultaneously duplicated.

#### 1.3.4 WGD bifans show a strong enrichment with respect to SSD bifans when the target genes interact at the protein level

A major finding of our analysis is the strong enrichment of WGD-derived bifans compared to SSD-derived ones when the duplicated target genes are involved in protein-protein interactions. Also in this case (TarBase and PrePPI networks) the global enrichment for both putative WGD and SSD pairs is strong, with *Z_Putative WGD_* = 34 and *Z_SSD_* = 49, but not very informative.

The enrichment is clearly visible in fig.8,A in which we plot the same Z-scores of fig.7,A but constrained only to this type of bifans. About three-quarters of the intragenic Ohno-miRNA pairs are involved in such PPI-bifans, while this is true for only about the 30% of the SSD pairs (fig.8,C). The distributions of Z-scores (fig.3,B) with the SSD-derived pairs show a significant difference. These results align with the hypothesis that WGD genes are preferentially retained when stoichiometric constraints are critical for the proteins they encode [25, 26]. This principle is particularly relevant in the context of WGD-derived miRNAs, which regulate genes involved in protein-protein interactions and complexes (see Discussion). In contrast, the formation of SSD bifans appears to follow a two-step process that does not inherently account for stoichiometric balance, increasing the likelihood of imbalance when such structures arise.

**Figure 8.**
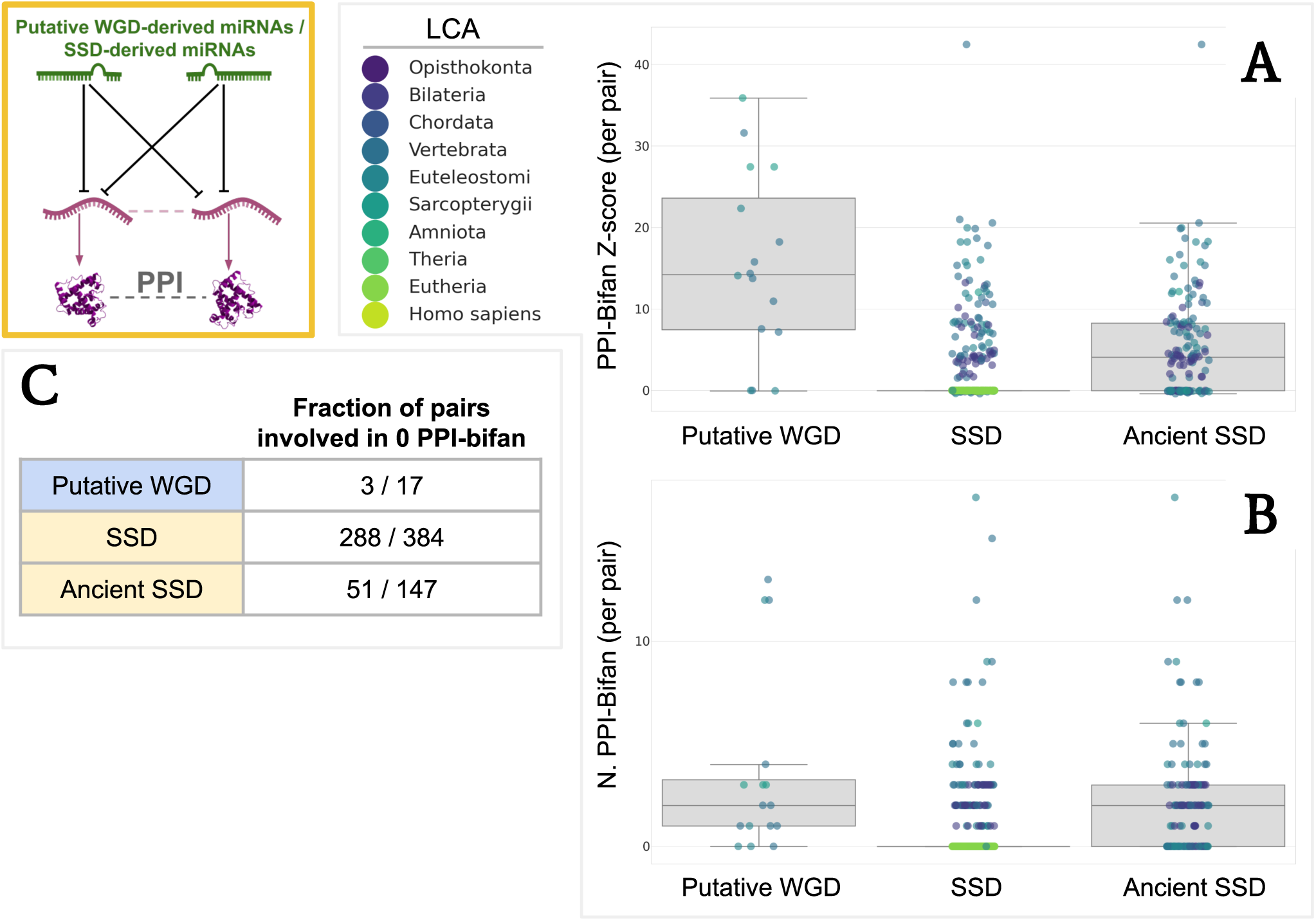
Same as fig.7 but for bifans in which the duplicated target genes interact at the protein level. Sub-figures, sets and color-coding are defined in the same way as explained in fig.7. P-values from the comparison of the Z-score distributions: *P_Putative WGD vs SSD_ <* 1*e−* 7, *P_Putative WGD vs Ancient SSD_ <* 1*e−* 3 (Kolmogorov-Smirnov test).

#### 1.3.5 PPI-Delta motifs

The so-called “delta motifs” are motifs in which two duplicated targets of the same miRNA are linked by a relation of interest, in our cases of two duplicated targets that also interact at the protein protein level. Here we show the enrichment of this motif for single miRNAs involved in WGD or SSD pairs (miRNA in WGD pairs that are also part of SSD pairs are excluded from the SSD sets, as described in Methods). Such delta motifs does not show a strong pattern differentiating single putative WGD and SSD miRNAs. The enrichment is negligible when analyzed from a global perspective with *Z_Putative WGD_* = 3 and *Z_SSD_* = 4 (the pairwise Z-score highlighted in Methods, fig.12,A can be trivially used to account for motifs involving single miRNAs like the delta motifs). Evaluating a Z-score for each miRNA belonging to a duplicated pair, Fig.9, shows how the few miRNAs with a high Z-score (Z>2) are exclusively part of SSD-derived pairs. The only perceptible difference in this analysis is a larger proportion of SSD miRNAs not involved in any delta motif, which however disappears when comparing WGD and Ancient SSD.

**Figure 9.**
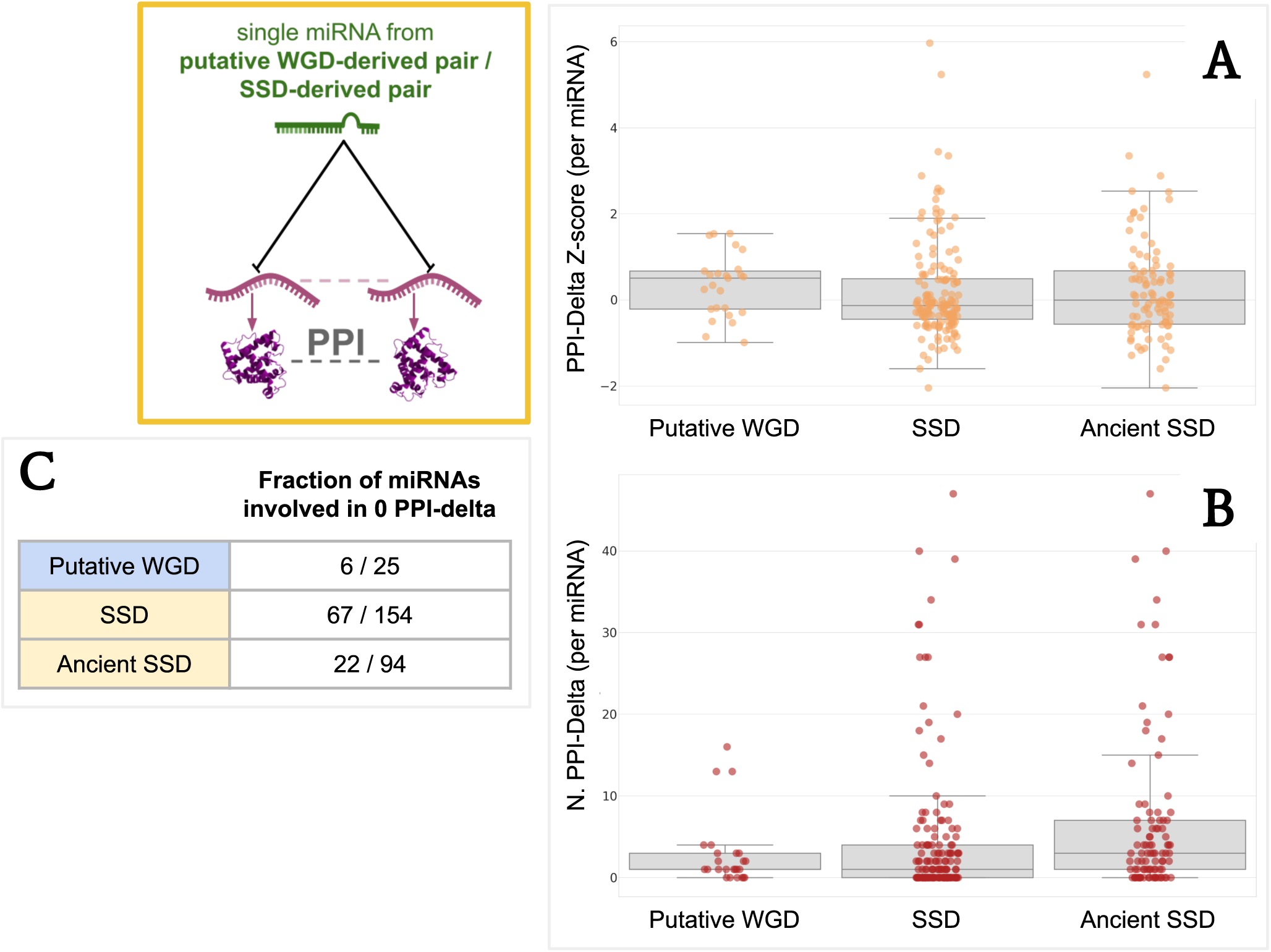
Enrichment of single miRNAs in delta motifs. The three sets are composed of individual miRNAs belonging to at least one pair of the same set as defined in Fig.7 and Fig.8. Individual miRNAs belonging to a WGD pair are discarded from both the SSD sets, even though they are present in SSD pairs. Color-coding is removed since a single miRNA can belong to different pairs with different last common ancestors. P-values from the comparison of the Z-score distributions: *P_Putative WGD vs SSD_ <* 0.01, *P_Putative WGD vs Ancient SSD_* = 0.11 (Kolmogorov-Smirnov test).

The near absence of any enrichment signal relative to these motifs suggests that the enrichment that we observed in fig.8 is linked to the particular topology of the PPI bifan and not to a generic preference of ohnolog miRNAs for duplicated pairs involved in protein-protein interactions. This signal is consistent changing the interaction network to MirDIP, and the protein-protein interaction network to STRING, with every possible combination (See Supplementary Material).

## 2 Discussion

### 2.1 A simple argument for the Ohno-miRNA retention

As reported in the Introduction, ohnolog genes are known to have many peculiar features and are involved in many crucial functions. These results are at odds with the expected backup role of duplicated genes (widely observed in less complex eukaryotes) which should provide a buffer against such effects. Within this context, the hypothesis of a prevalence of dosage-balanced genes among ohnologs has been proposed [5]. Changing the stoichiometry of members of a set of interacting genes (e.g. members of the same protein complex or the same pathway) may affect the function of the whole, resulting in detrimental effects on the fitness. This implies a selective pressure that keeps the balance between dosage of genes in these sets [27, 28]. WGD facilitates the maintenance of stoichiometric balance of all components of a dosage-balanced gene set compared to SSD. Moreover, if any gene is lost after a WGD it is likely to produce a dosage imbalance of the corresponding gene product, this may lead to preferential retention of dosage-balanced ohnologs [28]. Consistent with this assumption, retained ohnologs have been shown to be enriched for dosage-balanced genes that are resistant to subsequent SSD and to copy number variation in human populations [5].

A similar mechanism may explain the predisposition towards Ohno-miRNA retention within the human genome. It has been extensively established that a single miRNA typically targets the transcripts of hundreds of genes [29], effectively suppressing their translation [30]. Considering that these target genes are usually dispersed throughout the genome, WGD maintains the appropriate stoichiometric equilibrium between a miRNA and its entire target set, while SSD disrupts this equilibrium. This argument is even stronger for the intragenic miRNAs that we study in this paper, since they are known to be mostly co-transcribed with their host genes, (this is true in particular for the old intragenic miRNAs [31, 32] that we are addressing in this study) thus adding a constraint to the dosage balance scenario. In fact, whether a miRNA has its own promoter, thus having independent transcription, it could facilitate the modulation of its expression to avoid imbalance, but if it is co-transcribed then its modulation would also involve that of the host gene. It seems that essentially no intragenic miRNA (with a single exception that we shall discuss below) was retained after a small-scale duplication in the vertebrate lineage for more than 500 million years. A consequence of this reasoning is that the presence of a miRNA within a gene, even if it is not in dosage balance, reduces the probability that the gene can undergo a SSD fixed in the population, given that the SSD of the locus would also imply the duplication of the miRNA which is in dosage balance. Results resumed in figures 1 and 3 support this hypothesis, highlighting how SSD-derived gene pairs do not harbor pairs of intragenic SSD-derived miRNAs, with the exception of the peculiar case of MIR208A-MIR208B discussed in 2.5.

### 2.2 V-motif enrichment and regulatory redundancy

V-motifs are created straightforwardly by a duplication event, since immediately after the duplication the two miRNAs target the same set of genes. However, this redundancy is only temporary and in rather short evolutionary times the two miRNAs start to differentiate following the standard process of neofunctionalization or subfunctionalization which is the typical fate of duplicated genes [33, 34]).

This is clearly visible in fig.6,A where most of the SSD pairs have exactly zero V-motifs. Moreover, the fact that many of these events are recent (i.e., the last common ancestor is indicated in the Vertebrate clade or earlier clades, as shown in figure 6) suggests that the timescale of this evolutionary divergence is rather short and that even recently duplicated miRNAs have at present a negligible number of redundant regulatory interactions. On the contrary, the enrichment that we observe in those V-motifs involving Ohno-miRNAs suggests that for this class of duplicated miRNAs there is instead a strong evolutionary pressure to keep this regulatory redundancy. This is most likely a consequence of the dosage-balance constraint discussed above. This regulatory redundancy, previously observed at the transcriptional level in WGD-derived pairs of Transcription Factors [10], is confirmed at the post-transcriptional level. Regulatory redundancy is known to play an important role in shaping the complexity of the organism [23, 35, 36]. It increases the robustness of the network against mutations [37], it allows the implementation of complex regulatory mechanisms like the bifans that we shall discuss below. Moreover, redundancy, thanks to the different promoters in front of the two miRNAs allow their expression to be differentiated temporally or spatially [35]. This redundancy also enables independent tuning of their response to external stimuli [36], while maintaining regulation of the same target genes.

In this context, it’s crucial to notice that among the “Ancient SSD” miRNA pairs there are most probably several ohnolog miRNAs that we could not detect because of their intergenic location; this makes it interesting to compare these pairs isolated from the rest of recent SSD duplicates. The enrichment that we find for a few specific motifs should be considered as “lower bounds” of the actual enrichment. The very fact that, despite this bias, we find all the same relevant enrichment for these motifs suggests a special role played by the Ohno-miRNAs in the regulatory network.

### 2.3 Significance and functional roles of bifans in miRNA-target regulatory networks

The bifan is one of the most overrepresented network motifs in gene regulatory networks at the transcriptional level [38, 39]. Previous research on the importance of WGD duplications in the topology of regulatory networks highlighted that many motifs are particularly overrepresented when both the regulators and the targets are WGD or SSD genes [10]. We showed above that the same is true also at the post-transcriptional level. There are several reasons which may explain the specific importance of miRNA mediated bifans. They allow modulating in a finer way the expression of the targets, to differentiate (thanks to the different promoters in front of the two miRNAs) the response to external stimuli and more generally they can be considered as “decision-making” motifs[24]. These same arguments hold also for the transcriptional version of these motifs, what makes the miRNA-mediated version special is that miRNAs are known to act as fine-tuner of gene expression and are thus particularly effective (and thus strongly preserved during evolution) in performing the complex and delicate functions discussed above.

### 2.4 Synergy between protein-protein interaction and miRNA regulation

One of the most interesting outcomes of our analysis is that genes that interact at the protein level have a higher probability to be co-targeted by a pair of miRNAs in a bifan and this probability is even higher if both the miRNAs and the target are ohnolog genes. As we mentioned above, this could be explained by a dosage-balance mechanism driving WGD pair retention, but it is also an indication of the strong synergy between the protein-protein interaction layer and the post-transcriptional layer of regulation. This same synergistic behavior was observed in [10] at the transcriptional level. This is in line with the idea that miRNA regulation acts as a fine-tuning layer of regulation and is thus more immediately related to the stoichiometric constraints that are imposed by the presence of a protein-protein interaction between the two targets.

### 2.5 The case of MIR208A(MYH6)-MIR208B(MYH7)

The only pair of duplicated miRNAs hosted on a pair of SSD-derived genes is MIR208A (hosted on MYH6) - MIR208B (hosted on MYH7), it is interesting to address this case in more detail. The two host genes are long-known paralogue genes [40, 41] that are located close together on chromosome 14 (14q11.2). According to Ensembl, their last common ancestor was in the clade of *Opisthokonta* while MIR208A and MIR208B are recognized as paralogues by Ensembl but their last common ancestor is reported to be in the clade of *Euteleostomi*. Thus, the duplication of the two miRNAs should have occurred after the duplication of their host genes as well as after the two rounds of Vertebrate-specific WGDs, according to Ensembl. We consider this scenario to be very unlikely. We hypothesize that, after the WGD event that generated the pair composed of MIR499 and MIR208A-MIR208B ancestor with the respective host genes, a subsequent SSD duplication gave rise to the MIR208A-MIR208B pair, as outlined in fig. 10. It is interesting to notice that this pair is not independent of the Ohno-miRNA: since MYH7-MYH7B is an ohnolog gene pair and so it’s MYH6-MYH7B, MIR20B (MYH7) - MIR499A (MYH7B) and MIR208A (MYH6) - MIR499A (MYH7B) are found to be Ohno-miRNA by our analysis.

**Figure 10.**
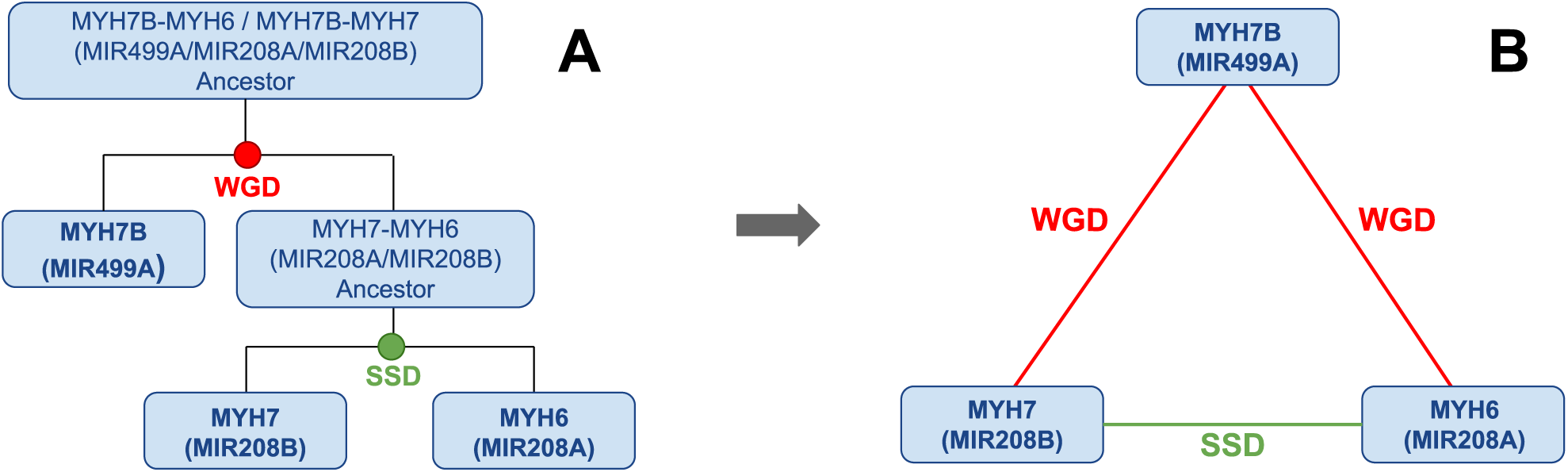
Suggested evolutive history linking MYH7B, MYH7 and MYH6 and their intronic miRNAs (MIR499A, MIR208A, MIR208B). As explained in the first paragraph of Section 2.5, the duplication times suggested by Ensembl are considered to be errors in the annotations.

The sequence similarity between the mature miRNAs is very high considering that these duplicates are not recent, the sequences of the 3’ mature miRNAs differ by three nucleotides outside the seed region, meaning that the seed is perfectly conserved both in the 5’ and the 3’ mature miRNAs. Similarly, the mature miRNA hsa-miR-499a-5p conserves a nearly identical seed than hsa-miR-208a-3p and hsa-miR-208b-3p (just 1 nucleotide is different considering the 7mer seed), the rest of the sequence is very different. Despite being considered as *bona fide* miRNAs by MirGeneDB, MIR208A and MIR208B are not present in the TarBase network, thus making motif analysis based on experimentally verified interactions impossible. MIR499A, MIR208A and MIR208 are part of a set of miRNAs known as “myomiRs” [42], some of which have already been extensively studied. All three of these miRNAs are known to be co-transcribed alongside their host genes [43] and in terms of spatial expression, miR-208b and miR-499 are found in both skeletal and cardiac muscle tissue, whereas miR-208a is uniquely expressed in the heart [44, 45]. Moreover, miR-208a and miR-208b are known to be chamber-specific, paralleling their host genes, since miR-208a is reported to be abundant in atrial myocardium, while miR-208b is reported to be preferentially expressed in left ventricles [46, 47]. The spatial differentiation in the expression of these two particular miRNAs can account for their peculiar conservation, allowing them to avoid deleterious effects due to stoichiometric imbalance. Some previous results in miRNA literature analyzed the expression divergence between duplicated miRNAs across human samples, without observing a pattern of increased divergence for the paralogue miRNAs [48].

### 2.6 Intergenic Ohno-miRNAs

Our pipeline is limited to intergenic miRNA. Since we have seen in the previous sections that OhnoMiRNAs show a well-defined pattern of motif enrichment in the regulatory network, it is tempting to use this pattern as a proxy to identify other pairs of intergenic Ohno-MiRNAs. It is tempting to use this pattern as a proxy to identify other pairs of Ohno-MiRNAs outside the introns. A perfect benchmark for this study is represented by two pairs: MIR181A1-MIR181A2 and MIR181B1-MIR181B2 which were shown to be the result of Whole Genome Duplication events (see [49] for further details and for a careful phylogenetic analysis of the family). Looking at them as they are reported as paralogues pairs in Ensembl, we found that they show high Z-scores for the network motifs which are typically enriched in Ohno-miRNAs pairs, as it’s visible from tab. 3.

**Table 3.**
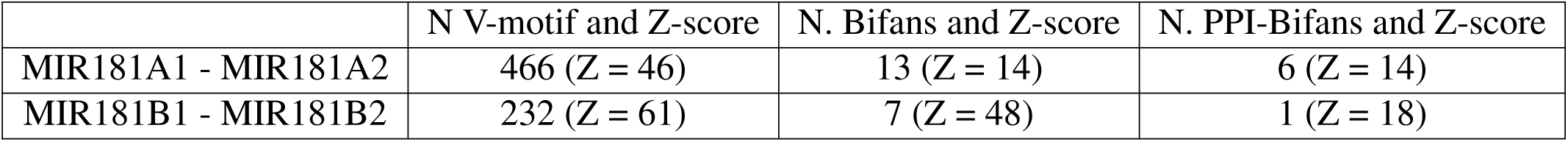
Number and Z-scores of different motifs involving the two MIR181 pairs. These results are obtained considering the TarBase network for the miRNA-target interactions and the PrePPI network for the protein-protein interactions. Z-scores for the bifans are computed considering ohnolog gene pairs as target.

|  | N V-motif and Z-score | N. Bifans and Z-score | N. PPI-Bifans and Z-score |
| --- | --- | --- | --- |
| MIR181A1 - MIR181A2 | 466 (Z = 46) | 13 (Z = 14) | 6 (Z = 14) |
| MIR181B1 - MIR181B2 | 232 (Z = 61) | 7 (Z = 48) | 1 (Z = 18) |

This is a non-trivial test of the whole analysis and we plan to use these considerations to extend our study also to intergenic miRNAs. A potential source of putative Ohno-miRNAs is the OHNOLOGS V2 database[6]. The intragenic pairs defined in our study are not included in this resource, likely due to challenges in defining orthology for non-coding genes. However, these challenges do not affect our pipeline, as we rely on the ohnology of the hosting protein-coding genes. We performed all the motif enrichment analysis on these pairs, tab.4 shows how most of them, especially among those corresponding to the *strict* criterion, show a strong enrichment in V-motifs, bifan and tends to be involved in PPI-Bifan with WGD-duplicated target genes.

**Table 4.** Number and Z-scores of different motifs involving miRNA pairs in the OHNOLOGS V2 database. Z-scores for the bifans are computed considering ohnolog gene pairs as targets. Different colors highlighting the pairs indicate different criteria on OHNOLOGS V2 (green=*strict*, blue=*intermediate*, red=*relaxed*).

|  | N V-motif and Z-score | N. Bifans and Z-score | N. PPI-Bifans and Z-score |
| --- | --- | --- | --- |
| MIRLET7A3 - MIRLET7E | 266 (Z = 26) | 5 (Z = 6) | 1 (Z = 3) |
| MIRLET7A3 - MIRLET7F2 | 462 (Z = 40) | 15 (Z = 13) | 6 (Z = 11) |
| MIRLET7F2-MIRLET7A2 | 457 (Z = 44) | 15 (Z = 14) | 6 (Z = 13) |
| MIRLET7A3 - MIRLET7A2 | 926 (Z = 65) | 45 (Z = 24) | 14 (Z = 17) |
| MIR124-1 - MIR124-2 | 170 (Z = 63) | 2 (Z = 32) | 2 (Z = 45) |
| MIRLET7A3 - MIRLET7A1 | 1,067 (Z = 73) | 61 (Z = 29) | 20 (Z = 21) |
| MIRLET7E - MIRLET7A2 | 262 (Z = 28) | 5 (Z = 7) | 1 (Z = 3) |
| MIRLET7A1 - MIRLET7A2 | 926 (Z = 64) | 287 (Z = 55) | 14 (Z = 17) |
| MIR124-1 - MIR124-3 | 170 (Z = 64) | 2 (Z = 24) | 2 (Z = 45) |
| MIRLET7C-MIRLET7A2 | 303 (Z = 33) | 6 (Z = 9) | 1 (Z = 4) |
| MIR133A1 - MIR133A2 | 119 (Z = 62) | 2 (Z = 37) | 1 (Z = NaN) |
| MIR133A1 - MIR133B | 0 (Z = 0) | 0 (Z = 0) | 0 (Z = 0) |
| MIRLET7F2-MIRLET7A1 | 462 (Z = 41) | 15 (Z = 13) | 6 (Z = 12) |
| MIR7-2 - MIR7-3 | 338 (Z = 62) | 2 (Z = 7) | 0 (Z = 0) |
| MIR7-1 - MIR7-2 | 380 (Z = 60) | 3 (Z = 6) | 1 (Z = 5) |
| MIR1-1 - MIR1-2 | 66 (Z = 62) | 0 (Z = 0) | 0 (Z = 0) |
| MIR206 - MIR1-2 | 0 (Z = 0) | 0 (Z = 0) | 0 (Z = 0) |
| MIRLET7A3 - MIRLET7C | 303 (Z = 33) | 6 (Z = 9) | 1 (Z = 3) |
| MIR7-1 - MIR7-3 | 338 (Z = 57) | 2 (Z = 5) | 0 (Z = 0) |
| MIRLET7E - MIRLET7C | 247 (Z = 47) | 5 (Z = 15) | 1 (Z = 9) |
| MIR196A2 - MIR96B | 137 (Z = 45) | 5 (Z = 42) | 1 (Z = 31) |
| MIR1-1 - MIR206 | 0 (Z = 0) | 0 (Z = 0) | 0 (Z = 0) |
| MIRLET7C-MIRLET7F2 | 292 (Z = 44) | 7 (Z = 17) | 1 (Z = 6) |
| MIRLET7C-MIRLET7A1 | 303 (Z = 33) | 6 (Z = 8) | 1 (Z = 3) |
| MIR9-1 - MIR9-3 | 482 (Z = 67) | 14 (Z = 25) | 2 (Z = 9) |
| MIR9-1 - MIR9-2 | 482 (Z = 68) | 14 (Z = 25) | 2 (Z = 9) |
| MIRLET7E - MIRLET7A1 | 266 (Z = 26) | 16 (Z = 7) | 1 (Z = 3) |

### 2.7 Robustness of the results

In the pipeline discussed in this paper we selected genes and interactions according to a few databases that represent the state-of-the-art in the field. In particular, miRNA-target interactions were described using the TarBase database and protein-protein interactions using the PrePPI database. It is interesting to see what happens to our results if we choose different databases for the interactions. We also evaluated the potential impact of literature bias in TarBase—arising from the overrepresentation of targets associated with extensively studied miRNAs—by comparing the results obtained from the entire database with those derived exclusively from high-throughput datasets. In particular, as discussed above, we chose as alternative databases MiRDIP for the miRNA-target interactions and STRING for the protein-protein interactions. As for those used in the main text, these are also state-of-the-art databases, but they select the interactions with criteria that are, so to speak, orthogonal with respect to the previous ones. In fact, as mentioned above, they show a very small number of common interactions. In the Supplementary Material we performed the same analyses discussed in the main text, changing the databases in all the possible ways. It turns out that our results, both for the miRNA selection and for the enrichment patterns are essentially the same, regardless of the choice of filter or database. We consider this as a major test of the robustness of our results.

## 3 Methods

### 3.1 SSD-derived and WGD-derived gene pairs

Following Mottes et al. [10] we obtained the WGD-derived pairs of ohnolog genes by merging the results of [5] with the list of human ohnologs available from the OHNOLOGS V2 database [6]. From the OHNOLOGS V2 database, we kept only the pairs corresponding to the *strict* criterion in order to filter high-confidence pairs of ohnologs. Only protein-coding genes were considered. The list of human protein-coding paralogue gene pairs was obtained from the Ensembl database [50]. We removed from this list all the pairs that were identified as WGD-derived. In order to ensure compatibility among different datasets, all the genes were traced back to their corresponding occurrences in the comprehensive gene annotation of the GENCODE database [51]. Following these criteria, we identified 9,348 WGD-derived pairs of protein-coding genes, involving 7,775 single genes, and 122,863 SSD-derived pairs of protein-coding genes involving 13,784 single genes. Considering single genes, 6,047 genes are involved both in a putative WGD-derived pair and in a putative SSD-derived pair.

### 3.2 Detection of Ohno-miRNA pairs

Among the gene pairs of our interest, those harboring a miRNA were detected using the BEDTools suite [52]. Given the list of WGD-derived and SSD-derived pairs, only pairs where both genes host a miRNA have been identified. We call an “Ohno-miRNA” pair, a pair of miRNA such that each miRNA is hosted on a protein-coding gene, and the two genes form a WGD-derived pair. We have only considered miRNAs recognized as *bona fide* by MirGeneDB [18]. A further check was performed on the strand of miRNAs with respect to the host genes in order to ensure that the relative orientation of the miRNA with respect to the hosting gene is the same for both the miRNAs in the pair. Given the case in which a gene belonging to a duplicate pair harbors more than a miRNA, our pipeline artificially inflates the number of putative Ohno-miRNA pairs providing every possible pair involving. In order to avoid this, given all the possible pairs formed by miRNAs harbored on a duplicated pair, only those pairs persisting after a reciprocal best alignment hit procedure were considered as putative Ohno-miRNA.

The study implemented a reciprocal best alignment hit approach, leveraging sequence similarity among miRNAs within gene pairs through the alignment method outlined in Section 3.3. For a given pair of ohnolog genes, each harboring multiple miRNAs, the pipeline considered all the possible miRNA pairs, pairing a miRNA from one ohnolog gene with every miRNA from the other. Designating *O*_1_ and *O*_2_ as generic ohnolog genes with multiple miRNAs, the analysis involved selecting each miRNA from *O*_1_ and identifying the most similar miRNA in *O*_2_ based on sequence similarity score, resulting in the best hit for each miRNA in *O*_1_. This procedure was reciprocally replicated for *O*_2_, yielding the best hit in *O*_1_ for every miRNA in *O*_2_. The final step of the reciprocal best alignment hit process entailed retaining pairs where the best hit was mutual, confirming a reciprocal match (i.e.„ *M*_1_ is the best hit of *M*_2_ and vice versa). The same procedures and sources have been used to retrieve putative intragenic ohnolog miRNAs in the mouse genome, described in Section 1.2.1, except for the list of protein-coding host genes. In fact, hnolog gene pairs in the mouse genome were obtained as the ortholog of the human protein-coding gene pairs. Orthology relations are obtained from the Ensembl database, keeping *one-to-many* relationships where more than an orthologue is present.

To further illustrate the reciprocal best alignment hit procedure, considering the ohnolog gene pair comprising DNM2 and DNM3: our analytical pipeline identified a solitary miRNA (MIR199A1) within DNM2, whereas DNM3 harbored two miRNAs (MIR199A2 and MIR214). After performing the reciprocal best alignment hit, the pair consisting of MIR214 and MIR199A1 was excluded, and only the MIR199A1-MIR199A2 pair was retained for further analysis.

### 3.3 Aligning miRNA sequences

The list of human miRNAs is retrieved from the GENCODE database. In order to obtain every mature miRNA originating from each miRNA gene we leveraged data from miRBase v22 [53] and Ensembl. Through miRBase, it was also possible to obtain the sequences for each mature miRNA and each pri-miRNA originating from a miRNA gene of interest. Given a putative Ohno-miRNA pair, we want to assign a similarity score based on the sequences. To do so, we retrieved every mature miRNA originating for a miRNA gene and aligned it with every other mature miRNA originating from the putative duplicated miRNA. The sequence similarity assigned to the Ohno-miRNA pair is the highest score obtained with this procedure.

The alignment score was established using a modified version of the Needleman-Wunsch algorithm, where the match and mismatch were assigned a greater (or lower in case of mismatches) weight in the substitution matrix, outlined in fig. 11. The seed was considered to be formed by nucleotides from 2 to 8 in a 7mer seed, starting numbering at the 5’ end of the mature miRNA. Matches and mismatches not involving nucleotides in the seed are given a weight of *±*1, while for in-seed matches and mismatches the weight is assigned to be *±* 5. The results are always consistent considering different lengths of the seed (*6mer*, *8mer*) and different weights assigned to in-seed matches and mismatches (*±*3, *±*4, *±*6).

**Figure 11.**
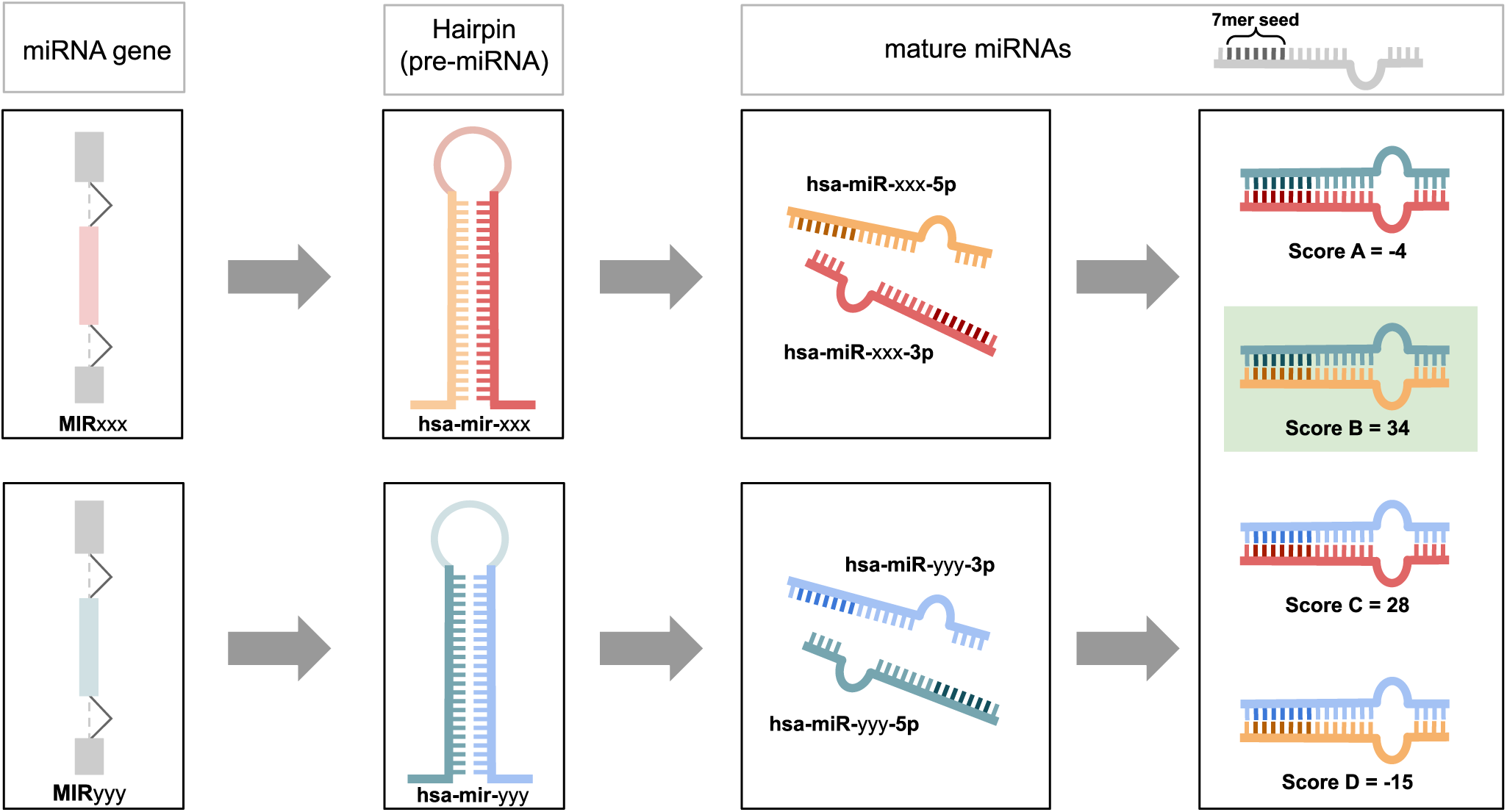
Schematic representation of how the alignment score is assigned to miRNA gene pairs, leveraging mature miRNA alignments. Mature miRNAs are aligned using the modified version of the Needleman-Wunsch described in Section 3.3. The represented case would see the pair MIRxxx and MIRyyy assigned an alignment score of 34.

### 3.4 SSD-derived miRNA pairs for the motif analysis

To compare the results obtained for Ohno-miRNAs, the identical procedures used to identify the Ohno-miRNAs were performed to retrieve similarly defined pairs of miRNAs harbored on SSD-derived gene pairs. This procedure precisely followed the steps delineated in fig.1. Removing from the obtained miRNA pairs those pairs that are not recognized as true paralogues by Ensembl, only two pairs survived: MIR208A(MYH6)-MIR208B(MYH7) and MIR196A1 (HOXB7) - MIR196A2 (HOXC6). The first pair is discussed in 2.5, while the second pair was an obvious example of misclassification, which we discuss in detail in the next section.

Results on the network motif analysis compare the list of putative intragenic ohnolog miRNA pairs (”putative WGD”) with a list of 685 duplicated miRNA pairs (involving 191 single miRNAs) from the Ensembl database (labeled “SSD” in the plots), filtered removing those recognized by our analysis as putative intragenic ohnolog and such that both miRNAs in a pair are *bona fide* miRNAs according to MirGeneDB. OHNOLOGS V2 database recognizes some pairs of WGD-derived miRNAs (28 pairs with the “relaxed” criterion, with one single overlap with our Ohno-miRNAs being MIR33A-MIR33B); see 2.6.

To minimize the number of likely spurious pairs among SSD miRNAs, we removed all pairs reported in OHNOLOGS V2 by adopting the “relaxed” filter from the analysis of V-motifs, bifans and PPI-bifans. From the analysis of single miRNAs in delta motifs, we also removed from the SSD sets all those miRNAs that are found to be involved only in pairs recognized as ohnolog by OHNOLOGS V2 and in no other pair in the Ensembl database.

Most of the plots also include a set of duplicated miRNA pairs labeled “Ancient SSD” which is a subset of the “SSD”-labeled pairs, obtained by keeping only pairs whose last common ancestor is comparable with *Vertebrata* (following [7], every pair whose last common ancestor is older than *Sarcopterygii*). Among the SSD-derived miRNA pairs, 174, involving 111 miRNAs, are labeled as “Ancient SSD”.

### 3.5 Manual classification of MIR196A1-MIR196A2

The pairing of MIR196A1 and MIR196A2 represents an obvious case of Ohno-miRNA misclassification as paralogues. Within the GENCODE database, two distinct annotation sets, namely *comprehensive* and *basic*, are available. The comprehensive annotations encompass a wide array of transcripts, including potentially rare or contextually specific ones. Conversely, the basic annotations emphasize a curated subset of transcripts, prioritizing those that are full-length and protein-coding. The overlap of MIR196A1 and MIR196A2 results with HOXB7 and HOXC6, respectively, is due to two respective non-basic transcripts characterized by extensive introns in UTR regions. Given that HOXB7 and HOXC6 are paralogues and not ohnologs, this miRNA pair is categorized as paralogue. However, considering their positioning within their respective HOX clusters, there is a compelling argument to reclassify them as Ohno-miRNAs. It is interesting to notice that studies referring to the miRNAs in the HOX clusters seem to have shown that MIR-196 has a known homologous (namely *mir-iab-4* in *Drosophila*) outside the vertebrate lineage [54] but is extensively present in numerous vertebrates, suggesting an origin and gaining of importance in the common ancestor of vertebrates which fits with the hypothesis of its origin after the whole-genome duplication. Therefore, we argue that these miRNAs should be more appropriately regarded as intergenic ohnologs rather than intronic paralogues. For these reasons, we manually reclassified MIR196A1 and MIR196A2 as Ohno-miRNAs prior to the motif analysis.

### 3.6 miRNA-target interaction networks

The miRNA-target networks used to assess motif enrichment and target similarity come from the TarBase [19] and the miRDIP [20] databases. TarBase is limited to experimentally validated interactions, while miRDIP integrates miRNA-target interactions coming from different databases and prediction methods. Both the networks are somehow biased, but in opposite directions. Literature-based collections like TarBase are characterized by numerous missing interactions (false negatives), moreover they are biased towards genes that received more attention from the scientific community. As pointed out in the Introduction, WGD-derived genes were shown to be often associated with diseases and organism complexity, which are preferential subjects of published papers. On the other hand networks based on in silico predictions of the interactions like MirDIP are in general characterized by a large number of false positives. Notwithstanding these differences, we shall see in the following that the two networks lead essentially to the same enrichment patterns (see Supplementary Material, Section 3). We consider this as a strong evidence of the overall robustness of our results.

The procedures used to obtain suitable networks from these databases are partially borrowed from [10]. The TarBase network was constructed by selecting all the interactions coming from Normal/Primary cell lines or tissues (excluding “cancer” and “other” categories), with positive evidence for a direct interaction between the miRNA and the target gene. Moreover, we only kept those interactions reported to be obtained with a high-throughput approach. The resulting network counts 713 miRNAs and 10,458 target genes, combined in 102,774 interactions. The miRDIP network was parsed leveraging the presence of an integrative score assigned by the authors to each miRNA-target pair. It is well established that the interactions provided by means of a miRNA-target prediction program are often noise (false positives or biologically irrelevant) [55], to overcome this problem, we decided to keep only the interactions belonging to the “Very high” score class (top 1% interactions in the database). In this case the resulting network is larger, and it’s composed of 1,847 miRNAs and 15,738 targets combined in 465,874 interactions. Since TarBase and miRDIP report data related to mature miRNA, each mature miRNA was traced back to the miRNA gene of origin in order to build a direct network involving miRNA genes instead of mature miRNAs (see 3.8). The resulting networks are very similar in terms of nodes, as 713 miRNAs and 9,161 genes are in common between TarBase and MirDIP, but different in terms of the interactions as only 9,809 edges are in common between the two networks.

A short description of the two networks is reported in Section 6 of the Supplementary Material.

### 3.7 Protein-protein interaction networks

Consistent with [10] protein-protein interactions were retrieved from two distinct databases: PrePPI [21] and STRING [22]. From PrePPI we kept only the high-confidence interactions with an experimental validation from the HINT database or in the APID database. The STRING database was parsed, selecting a high confidence score (>960) to keep the size of the two PPI-networks comparable. The protein IDs were mapped to the corresponding gene name using the UniProtKB Database [56].

We ended up with 45,386 PPIs and 8,944 genes in the PrePPI network, while the resulting STRING network is composed of 51,268 PPIs and 9,758 genes. The two networks largely overlap in terms of nodes (6,730 genes are present in both networks) but are different in terms of interactions, with only 16,044 interactions in common. As for the miRNA-target interaction network, we provide a summary of the presence of ohnologs and paralogues in the two employed networks in *Section 6* of the Supplementary Material.

### 3.8 Sequence identity between duplicated miRNAs requires manual curation of the miRNA-target networks

When looking at enrichment scores, a potential problem of the TarBase and MirDIP databases is the fact that in some cases, duplicated miRNA genes can result in identical mature miRNAs after the processes of transcription and cleavage and are treated by the databases as a single miRNA. Some examples of this redundancy can be found in the following miRNA pairs: MIR196A1 and MIR196A2 genes are both transcribed and cleaved into the same 5’ mature miRNA named hsa-miR-196a-5p [57]. A similar case is represented by the pair MIR218-1 and MIR218-2, they have different 3’ mature miRNAs (hsa-miR-218-1-3p and hsa-miR-218-2-3p) but an identical 5’ mature miRNA (hsa-miR-218-5p). To address this issue, we manually curated the regulatory networks constructed from the two databases to keep the distinction between the miRNAs that were lost in the databases. In particular, we mapped each mature miRNA back to its miRNA gene, thus mapping the original “mature miRNA-mRNA” network in a “miRNA gene-mRNA” network. Whenever a mature miRNA originates from more than a miRNA gene, the new network has two different corresponding nodes.

### 3.9 Sørensen-Dice similarity coefficient

As a metrics for the interaction similarity of two duplicated miRNAs we used the Sørensen-Dice similarity coefficient, defined in the following way:

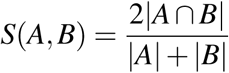

Where *A* and *B* are the sets of target genes of the two miRNAs in the pair. The Sørensen-Dice similarity coefficient is equal to 0 when there are no target genes in common between the two Ohno-miRNAs, and it’s equal to 1 when all interactions are in common.

### 3.10 Null models

To assess motif enrichment, we introduced an ensemble of 1,000 null models by applying random rewiring to our networks. We employed a degree-preserving procedure for randomization (as in [10]), preserving degrees for each node while randomly rewiring interactions between miRNAs and target genes. This ensured that every gene and miRNA maintained the same number of interactions as in the original network. By doing so, we aimed to investigate whether the observed enrichment patterns were influenced by degree-degree correlations within the duplicated pairs, as these correlations remained consistent in the randomized network set.

### 3.11 Quantifying ohnolog miRNA enrichment in different network motifs

Network motifs refer to specific patterns of nodes and edges that exhibit a notable overrepresentation in the regulatory network when compared to randomized networks [58]. It is widely accepted that these motifs have undergone positive evolutionary selection owing to their functional efficacy. To evaluate the enrichment of network motifs, we present the Z-score associated with the motif count within the ensemble of randomized networks. The Z-score calculation is generally given by:

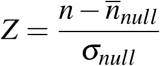

where *n* is the motif count on the real network, *n_null_*is the average motif count computed across 1,000 randomized networks, and *σ_null_* is the standard deviation derived from the same distribution. We present the Z-scores computed on the single pair (or single miRNA in the case of PPI-delta). Differences with the standard way to evaluate the Z-scores, where *n* is the number of motifs counted over all the pairs in a set without caring how many motifs involve which nodes, are depicted in Fig. 12. We plotted only the Z-score of putative intragenic ohnolog miRNA pairs recognized as actual duplicated pairs by Ensemble, since the set of SSD duplicated pairs are downloaded from the Ensembl. If *n_null_* = 0 and *σ_null_* = 0 (i.e., at least one motif is present in the real network but we couldn’t find a motif involving the pair in any of the 1,000 randomized networks) the pair is discarded because it would be impossible to evaluate the Z-score. On the other hand, if *n_null_*= 0 and *σ_null_*= 0, we set *Z* = 0. Discarded pairs never reach more than the 1% of the total pairs in a given set. However, since these cases involve pairs relevant to the analysis, we report the discarded pairs in the Supplementary Material. Pairs where one or both miRNAs were not present in the interaction network were obviously discarded. When analyzing delta motifs (Section 1.3.5) individual miRNAs belong to both WGD and SSD pairs: for example MIR10A is in a WGD pair with MIR10B according to our analysis, and in a SSD pair with MIR125A according to Ensembl. In this case these miRNAs would be both in the “putative WGD” and in the SSD sets. In order to avoid this, if a miRNA is part of a putative WGD pair it is removed from the SSD, and possibly Ancient SSD, set.

**Figure 12.**
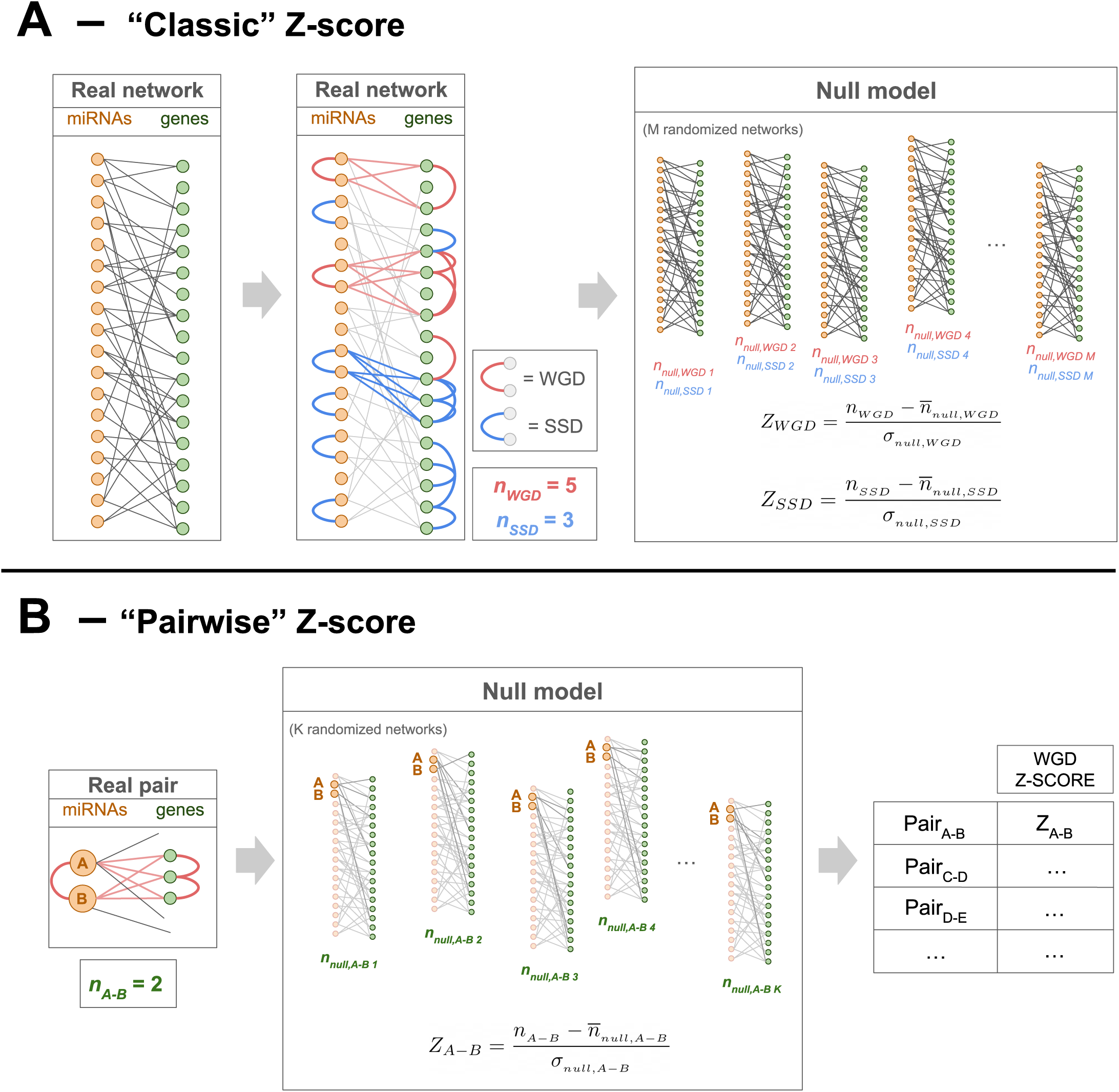
Differences between the “classic” and a “pairwise” way of reporting the Z-scores for subset of nodes in a given network (the example considers bifans). In case **(A)** we sum the counts from all the pairs belonging to the same set (e.g. SSD or WGD), thus not knowing if the resulting Z-score is due to all the pairs in the set or just to a subset. On the other hand, in case **(B)** we count the number of motif and generate a distribution of counts over the null model for each pair in each set, obtaining a distribution of Z-scores for each set (WGD, SSD and Ancient SSD). Pairwise Z-score can be straightforwardly extended to single miRNAs when analyzing delta motifs.

## Concluding Remarks

The main goal of this paper was to identify a set of miRNA pairs derived from the two rounds of WGD events at the beginning of the vertebrate lineage and study their role in the regulatory network.

We identified a set of overrepresented regulatory motifs involving these miRNAs whose specific enrichment is likely to be due to the dosage balance constraint. We also realized that these Ohno-miRNA pairs show a strong tendency to keep the same seed sequence and to regulate common target genes. The combination of these two trends leads to an increase of redundancy of the regulatory network which is a typical hallmark of complexity [23]. Indeed, the same pattern was observed looking at WGD pairs of Transcription Factors and their enriched motifs in the transcriptional layer of the regulatory network [10]. Our results could be extended in two directions. First, the same approach used in this paper could be used to study other non-coding genes hosted in the introns of ohnolog protein-coding genes. In particular, it would be interesting to identify in this way WGD derived lncRNA and thus prioritize their study. Second, our analysis could be extended also to intergenic regions using the properties of the Ohno-miRNAs that we discussed in text as features of a classification algorithm combined with a systematic analysis of synthenic regions of WGD origin in the genome along the lines discussed in [6, 7] for the coding genes. Both these directions are worthwhile to be explored to further support the observation, which is the main result of our work, that the two rounds of Whole Genome Duplications played an essential role in increasing the complexity of the regulatory network, which is most likely at the origin of the impressive variety and complexity of the organisms belonging to the vertebrate lineage.

## Acknowledgements (not compulsory)

Leonardo Agasso is a Ph.D. student in Complex Systems for Quantitative Biomedicine, XXXIX cycle, at the University of Turin. We thank Matteo Osella, Francesco Mottes and Hervé Isambert for the useful discussions.

## Author contributions statement

M.C. and I.M. conceived and supervised the study; L.A., I.M., and M.C. identified key results. L.A. collected and processed the datasets, developed the methodologies and analyzed the data. All the authors wrote the manuscript and approved its final draft.

## Availability of data and materials

The makefiles, scripts and notebooks used to obtain every result presented in this paper will be publicly available at the GitHub repository https://github.com/LeonardoAgasso/OhnomiRNAs upon publication.

## Footnotes

1 The transcriptional version of this motif (in which a pair of paralogue transcription factors regulate a pair of paralogue targets) was studied in detail in [10]

