## Supplementary Material for "Ohno-miRNAs: intragenic miRNA pairs derived from whole-genome duplication"

### 1 Duplicated pairs before Ensembl paralogues filter

In Section 1 of the Main Text, we identified a list of putative duplicated pairs but we kept it for further analysis only those that are recognized as paralogues by Ensembl. Here we report the list of all the duplicated pairs.

**Table 1.** Putative ohnolog miRNAs with their respective host genes. Rows highlighted in red are those recognized as duplicated by the Ensembl database, and thus reported in the Main Text.

| Putative Ohno-miRNA 1 | Host gene 1 | Putative Ohno-miRNA 2 | Host gene 2 |
|---|---|---|---|
| MIR199A1 | DNM2 | MIR199B | DNM1 |
| MIR199A1 | DNM2 | MIR199A2 | DNM3 |
| MIR199B | DNM1 | MIR199A2 | DNM3 |
| MIR103A1 | PANK3 | MIR103A2 | PANK2 |
| MIR103A1 | PANK3 | MIR107 | PANK1 |
| MIR103A2 | PANK2 | MIR107 | PANK1 |
| MIR26B | CTDSP1 | MIR26A1 | CTDSPL |
| MIR26B | CTDSP1 | MIR26A2 | CTDSP2 |
| MIR26A1 | CTDSPL | MIR26A2 | CTDSP2 |
| MIR196A2 | HOXC6 | MIR196A1 | HOXB7 |
| MIR10A | HOXB3 | MIR10B | HOXD3 |
| MIR218-1 | SLIT2 | MIR218-2 | SLIT3 |
| MIR204 | TRPM3 | MIR211 | TRPM1 |
| MIR152 | COPZ2 | MIR148B | COPZ1 |
| MIR128-1 | R3HDM1 | MIR128-2 | ARPP21 |
| MIR153-1 | PTPRN | MIR153-2 | PTPRN2 |
| MIR33B | SREBF1 | MIR33A | SREBF2 |
| MIR499A | MYH7B | MIR208B | MYH7 |
| MIR499A | MYH7B | MIR208A | MYH6 |
| MIR3613 | TRIM13 | MIR16-2 | TRIM59-IFT80 |

**Table 2.** Putative SSD-derived miRNA pairs. Rows highlighted in red are those recognized as duplicated by the Ensembl database, and thus reported in the Main Text.

| Putative paralogue miRNA 1 | Host gene 1 | Putative paralogue miRNA 2 | Host gene 2 |
|---|---|---|---|
| MIR208B | MYH7 | MIR208A | MYH6 |
| MIR489 | CALCR | MIR642A | GIPR |
| MIR105-2 | GABRA3 | MIR452 | GABRE |
| MIR105-1 | GABRA3 | MIR452 | GABRE |
| MIR499A | MYH7B | MIR1266 | MYO5C |
| MIR499A | MYH7B | MIR3136 | TMF1 |
| MIR499A | MYH7B | MIR2116 | MYO1E |
| MIRLETF2 | HUWE1 | MIR140 | WWP2 |
| MIR208B | MYH7 | MIR1266 | MYO5C |
| MIR208B | MYH7 | MIR3136 | TMF1 |
| MIR208B | MYH7 | MIR2116 | MYO1E |
| MIR208B | MYH7 | MIR208A | MYH6 |
| MIR582 | PDE4D | MIR139 | PDE2A |
| MIR10A | HOXB3 | MIR615 | HOXC5 |
| MIR10A | HOXB3 | MIR196A2 | HOXC6 |
| MIR10A | HOXB3 | MIR615 | HOXC4 |
| MIR10A | HOXB3 | MIR196A1 | HOXB7 |
| MIR10B | HOXD3 | MIR615 | HOXC5 |
| MIR10B | HOXD3 | MIR196A2 | HOXC6 |
| MIR10B | HOXD3 | MIR615 | HOXC4 |
| MIR10B | HOXD3 | MIR196A1 | HOXB7 |
| MIR1266 | MYO5C | MIR3136 | TMF1 |
| MIR1266 | MYO5C | MIR2116 | MYO1E |
| MIR1266 | MYO5C | MIR208A | MYH6 |
| MIR561 | GULP1 | MIR556 | NOS1AP |
| MIR3136 | TMF1 | MIR2116 | MYO1E |
| MIR3136 | TMF1 | MIR208A | MYH6 |
| MIR218-1 | SLIT2 | MIR876 | LINGO2 |
| MIR589 | FBXL18 | MIR887 | FBXL7 |
| MIR2116 | MYO1E | MIR208A | MYH6 |
| MIR676 | EDA | MIR455 | COL27A1 |
| MIR101-1 | JAK1 | MIR151A | PTK2 |
| MIR615 | HOXC5 | MIR196A2 | HOXC6 |
| MIR615 | HOXC5 | MIR196A1 | HOXB7 |
| MIR876 | LINGO2 | MIR218-2 | SLIT3 |
| MIR196A2 | HOXC6 | MIR615 | HOXC4 |
| MIR615 | HOXC4 | MIR196A1 | HOXB7 |

#### 2 miRNA statistics: intragenic and intergenic miRNAs

In the human genome, miRNAs can be intragenic (hosted on a gene, usually on the introns) or intergenic. Here we summarize the nature of all the miRNAs in GENCODE. A miRNA is labeled as intragenic if it can be found in the intron (or exon) of at least a protein-coding gene transcript.

**Table 3.** Intragenic and intergenic miRNAs among the 1,877 miRNAs in the GENCODE database

| N. intragenic miRNAs | N. intergenic miRNAs |
|---|---|
| 1,200 (64%) | 677 (36%) |

**Table 4.** Intragenic and intergenic miRNAs among the 505 miRNAs in the GENCODE database that are recognized as *bona fide* by MirGeneDB

| N. intragenic miRNAs | N. intergenic miRNAs |
|---|---|
| 245 (49%) | 260 (51%) |

#### 3 MirDIP network

All the analyses in the Main Text conducted on the TarBase network have been replicated on the MirDIP network, keeping only the interaction in the *Very High* score class (which represent the top 1% interactions). Results obtained using the MirDIP network do not significantly differ from those presented in the Main Text.

##### 3.1 Target similarity

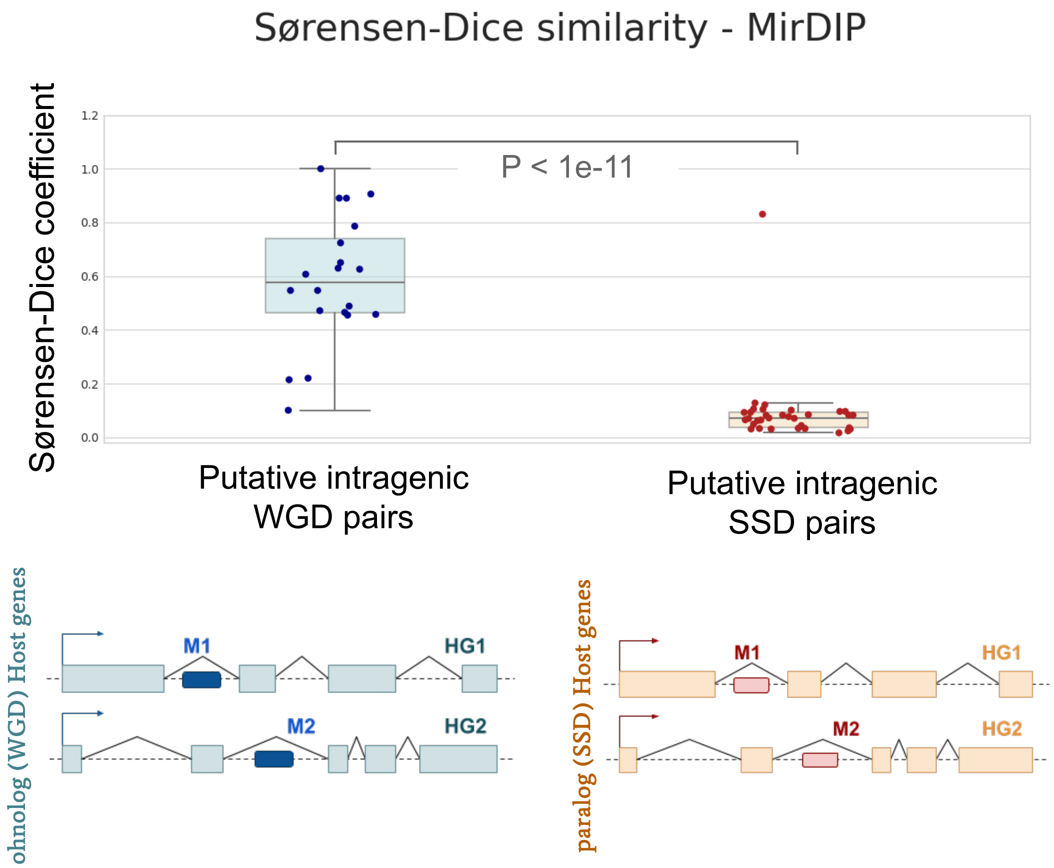


**Figure 1.** Distribution of the Sorensen-Dice coefficient of the duplicated pairs. This figure is the analogous of Fig.3,B, Main Text. Results are consistent with those obtained on the TarBase network.

#### 3.2 Outdegree distribution

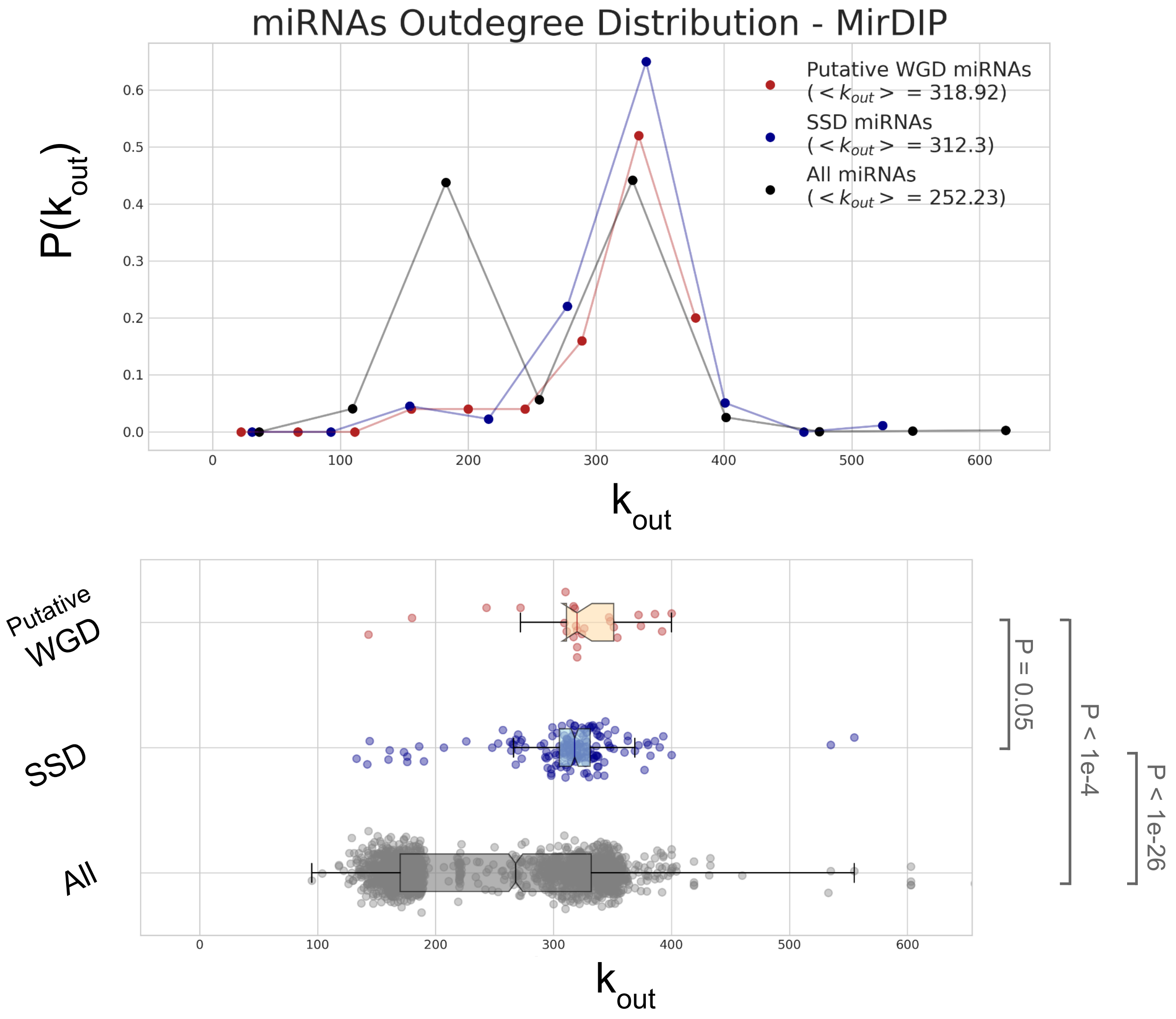


**Figure 2.** Out-degree distribution of putative Ohno-miRNAs compared with that of the paralogue miRNAs from Ensembl, and all the miRNAs in the MirDIP network.

##### 3.3 Simple motif enrichment: V-motif and bifan

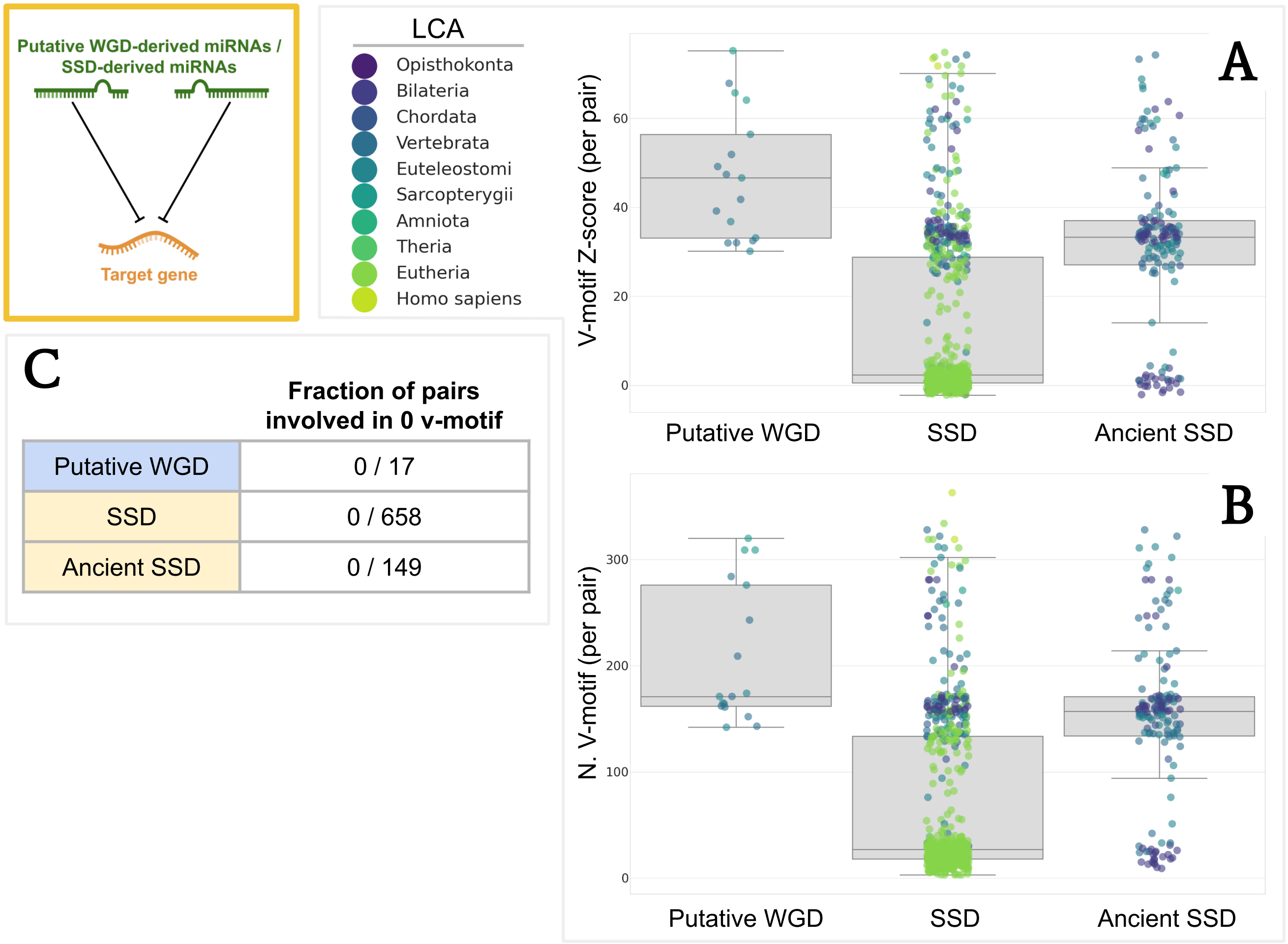


**Figure 3.** V-motif enrichment in the MirDIP network, this figure is the analogous of Fig.5 in the Main Text. **(A)** Distribution of the Z-scores refers to the enrichment of each pair with respect to the null model (each dot is a pair, the Last Common Ancestor of the pair is color coded according to the legend). The first set reports the z-score of motif enrichment of our putative Ohno-miRNA pairs, in the second and third set the same quantity for all the other pairs of paralogue miRNAs in Ensembl, in the third set (Ancient SSD) we keep only pairs whose Last Common Ancestor older than *Sarcopterygii*. **(B)** Number of V-motif involving each pair. **(C)** Number of pair not involved in any V-motif for each of the three sets.
P-values from the comparison of the Z-score distributions: $P_{PutativeWGD\ vs\ SSD} < 1e-10$, $P_{PutativeWGD\ vs\ Ancient\ SSD} < 0.01$ (Kolmogorov-Smirnov test).

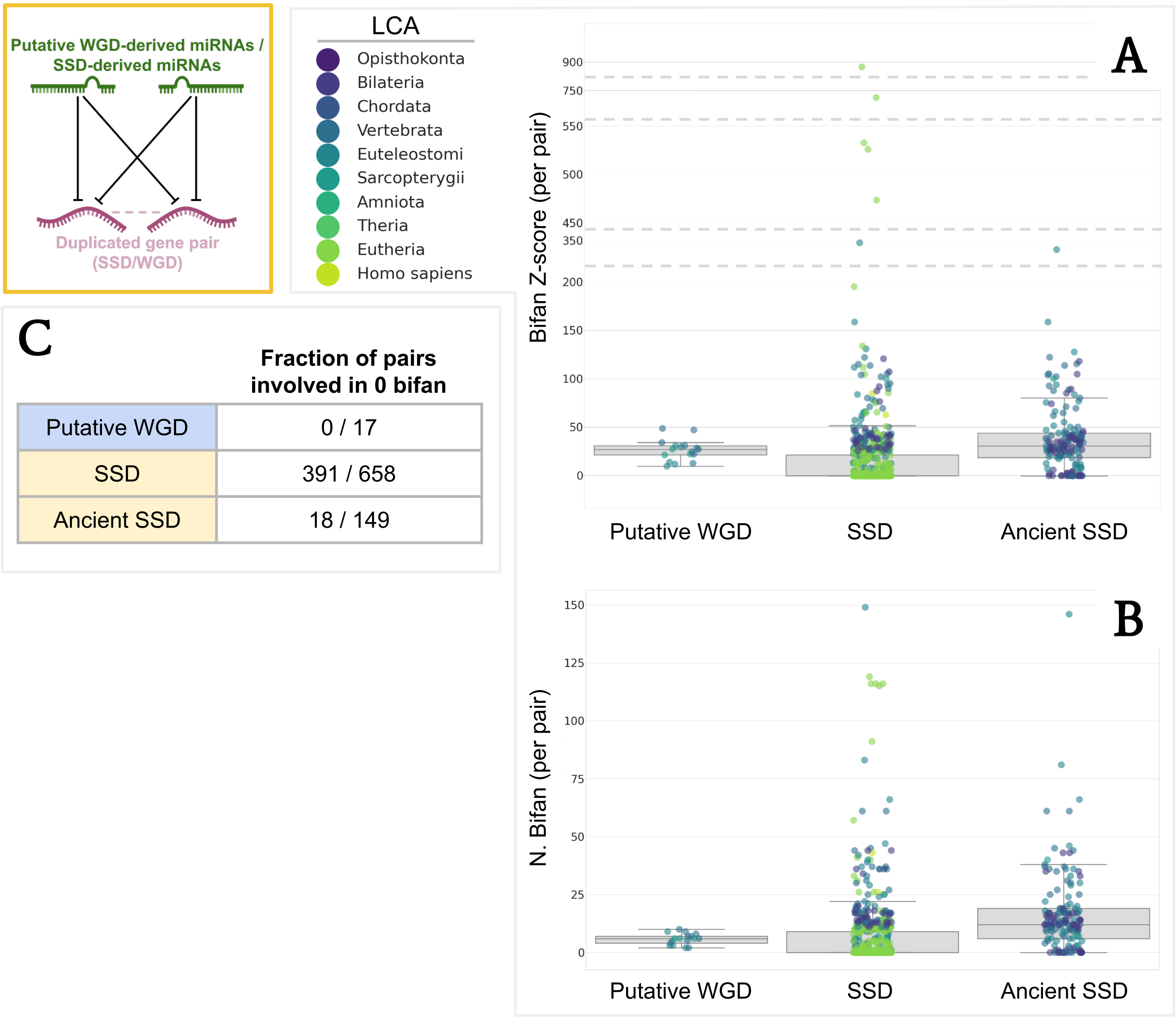


**Figure 4.** Bifan enrichment in the MirDIP network, this figure is the analogous of Fig.6 in the Main Text. **(A)** Distribution of the Z-scores referred to the enrichment of each pair with respect to the null model, the definition of the three sets and the color-coding is the same as ing Fig.3. **(B)** Number of V-motif involving each pair. **(C)** Number of pair not involved in any bifan for each of the three sets.
P-values from the comparison of the Z-score distributions: $P_{PutativeWGD\ vs\ SSD} < 1e-8$,
$P_{PutativeWGD\ vs\ Ancient\ SSD} = 0.23$

.

##### 3.3.1 MirDIP interaction network - PrePPI PPI network

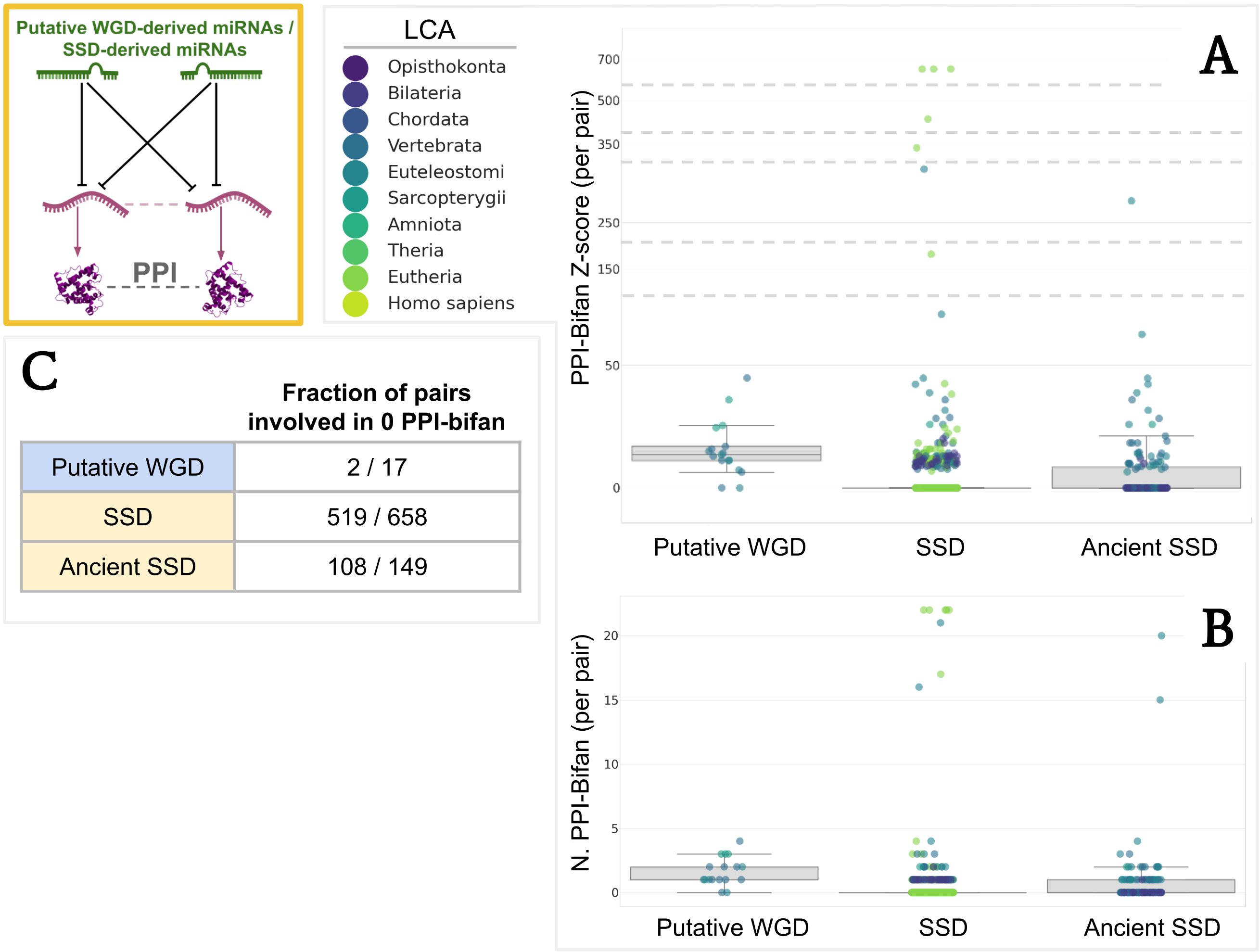


**Figure 5.** Same as Fig.6 of the Main Text, but changing the miRNA-target interaction network to MirDIP. P-values from the comparison of the Z-score distributions: $P_{PutativeWGD\ vs\ SSD} < 1e-7$, $P_{PutativeWGD\ vs\ Ancient\ SSD} < 1e-5$ (Kolmogorov-Smirnov test).

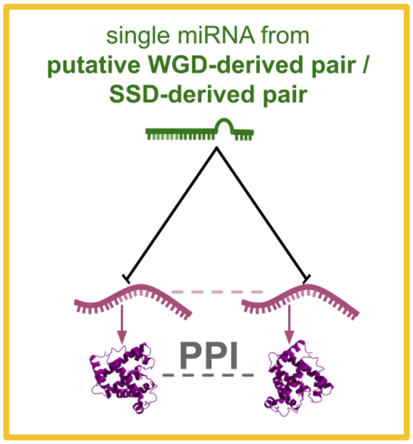


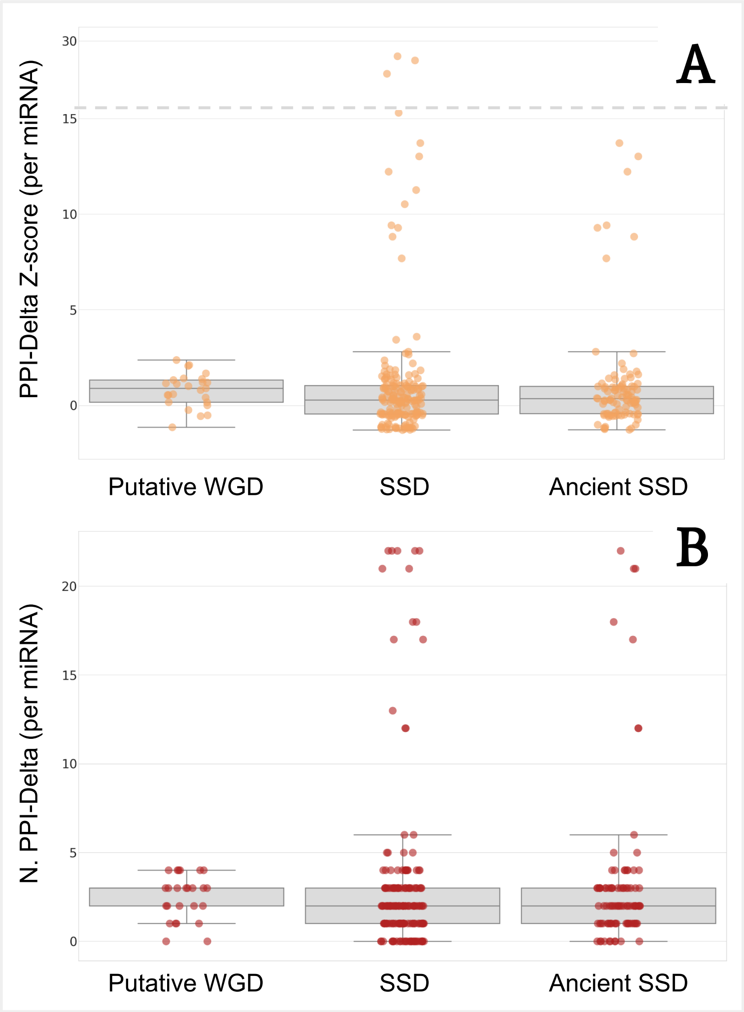


**C**

| | Fraction of miRNAs involved in 0 PPI-delta |
|---|---|
| Putative WGD | 2 / 25 |
| SSD | 30 / 177 |
| Ancient SSD | 10 / 97 |

**Figure 6.** Same as Fig.5 of the Main Text, but changing the miRNA-target interaction network to MirDIP. P-values from the comparison of the Z-score distributions: $P_{PutativeWGD\ vs\ SSD} = 0.03$, $P_{PutativeWGD\ vs\ Ancient\ SSD} = 0.07$ (Kolmogorov-Smirnov test).

#### 4 PPI motif enrichment - STRING

The results observed in Section 1.3 of the Main Text and Section 2.3.1 of the Supplementary can be tested by changing the protein-protein interaction network to STRING (see Section 3.7 of the Methods in the Main Text).

#### 4.1 TarBase interaction network - STRING PPI network

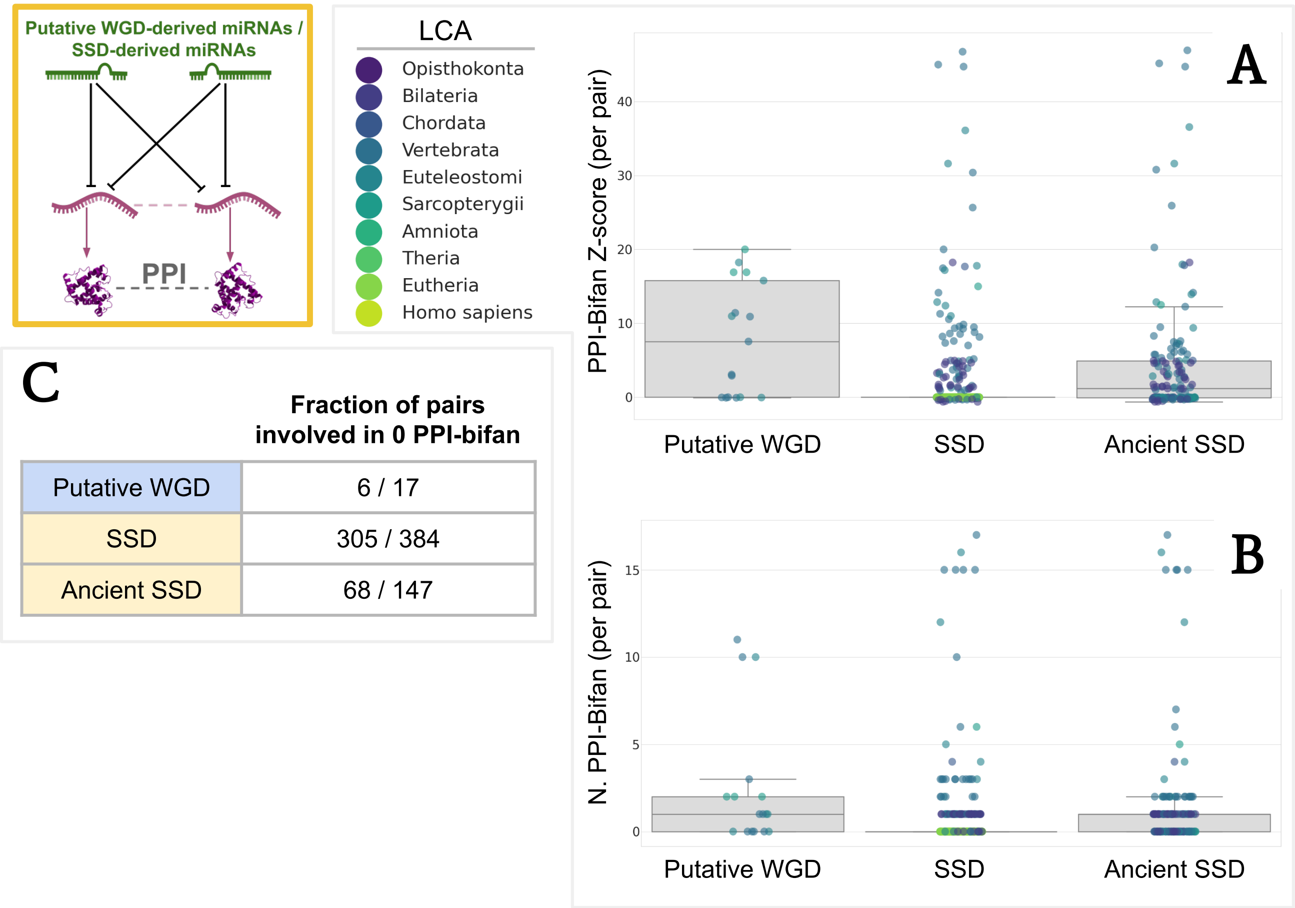


| | Fraction of pairs involved in 0 PPI-bifan |
|---|---|
| Putative WGD | 6 / 17 |
| SSD | 305 / 384 |
| Ancient SSD | 68 / 147 |

**Figure 7.** Same as Fig.7 of the Main Text but changing the protein-protein interaction network to STRING.
P-values from the comparison of the Z-score distributions: $P_{PutativeWGD\ vs\ SSD} < 1e-3$, $P_{PutativeWGD\ vs\ Ancient\ SSD} < 0.01$ (Kolmogorov-Smirnov test).

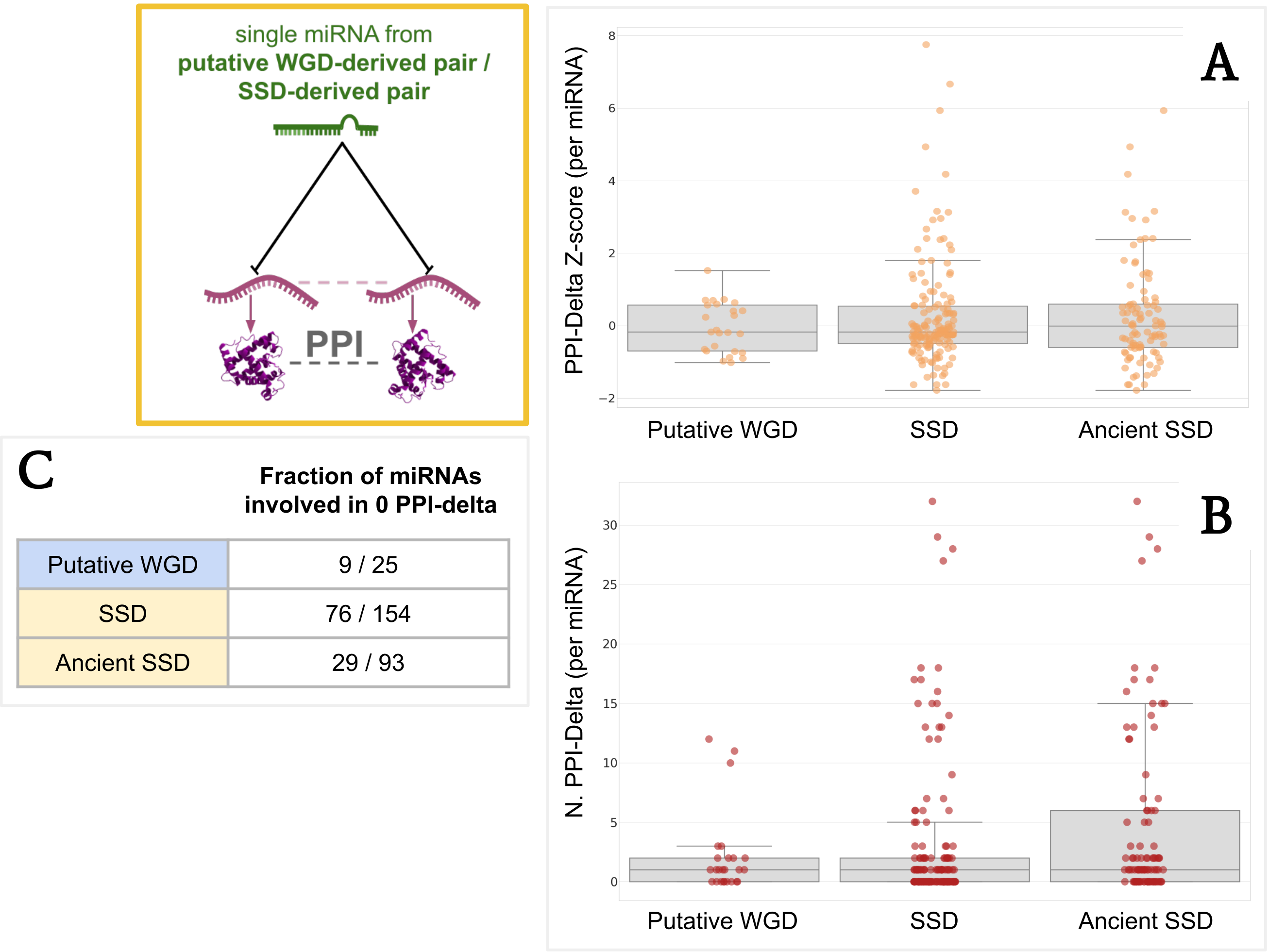


**Figure 8.** Same as Fig.8 of the Main Text but changing the protein-protein interaction network to STRING.
P-values from the comparison of the Z-score distributions: $P_{PutativeWGD\ vs\ SSD} = 0.49$, $P_{PutativeWGD\ vs\ Ancient\ SSD} = 0.48$ (Kolmogorov-Smirnov test).

##### 4.1.1 MirDIP interaction network - STRING PPI network

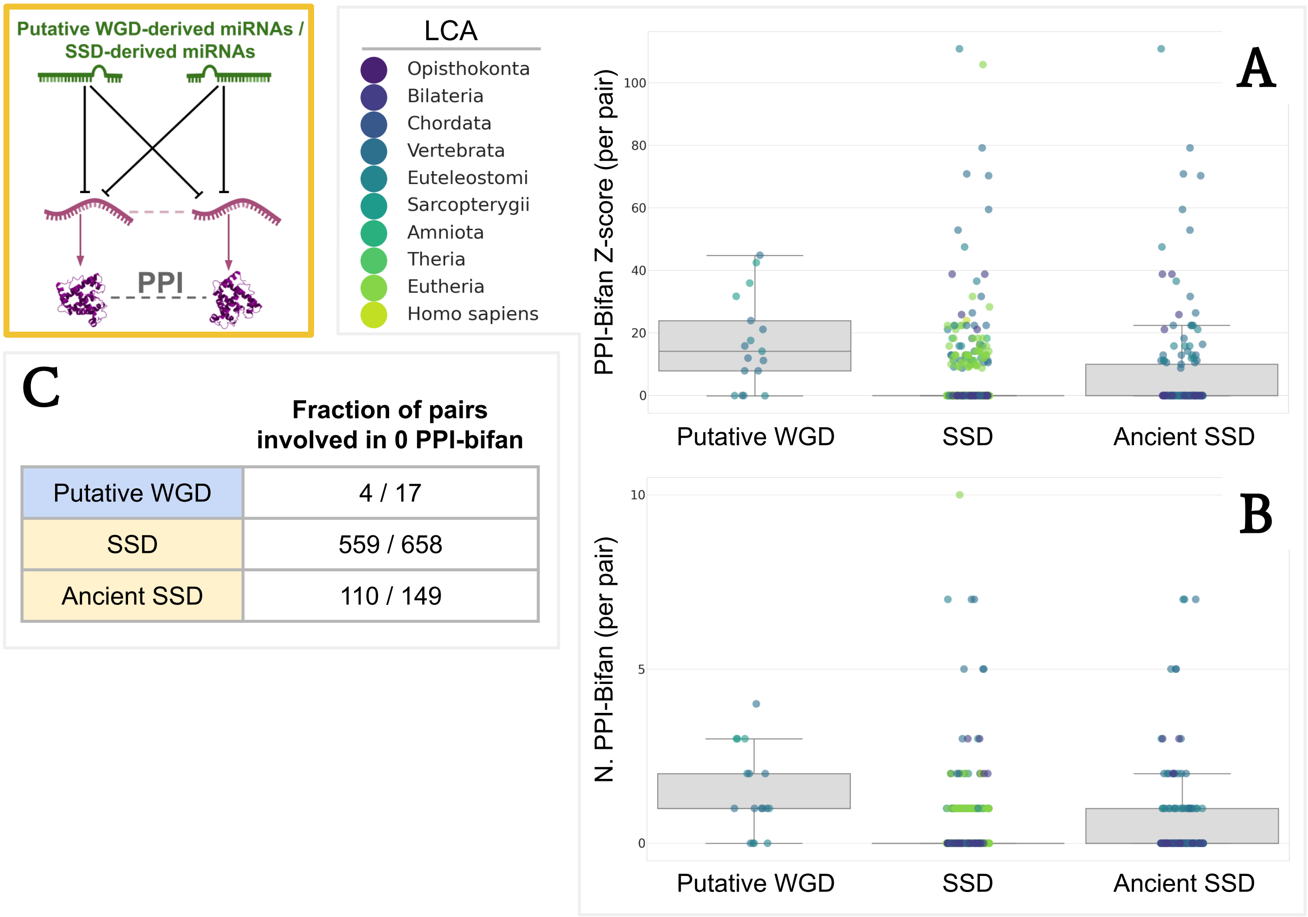


**Figure 9.** Same as 5 but changing the protein-protein interaction network to STRING. P-values from the comparison of the Z-score distributions: $P_{PutativeWGD\ vs\ SSD} < 1e-6$, $P_{PutativeWGD\ vs\ Ancient\ SSD} < 1e-4$ (Kolmogorov-Smirnov test).

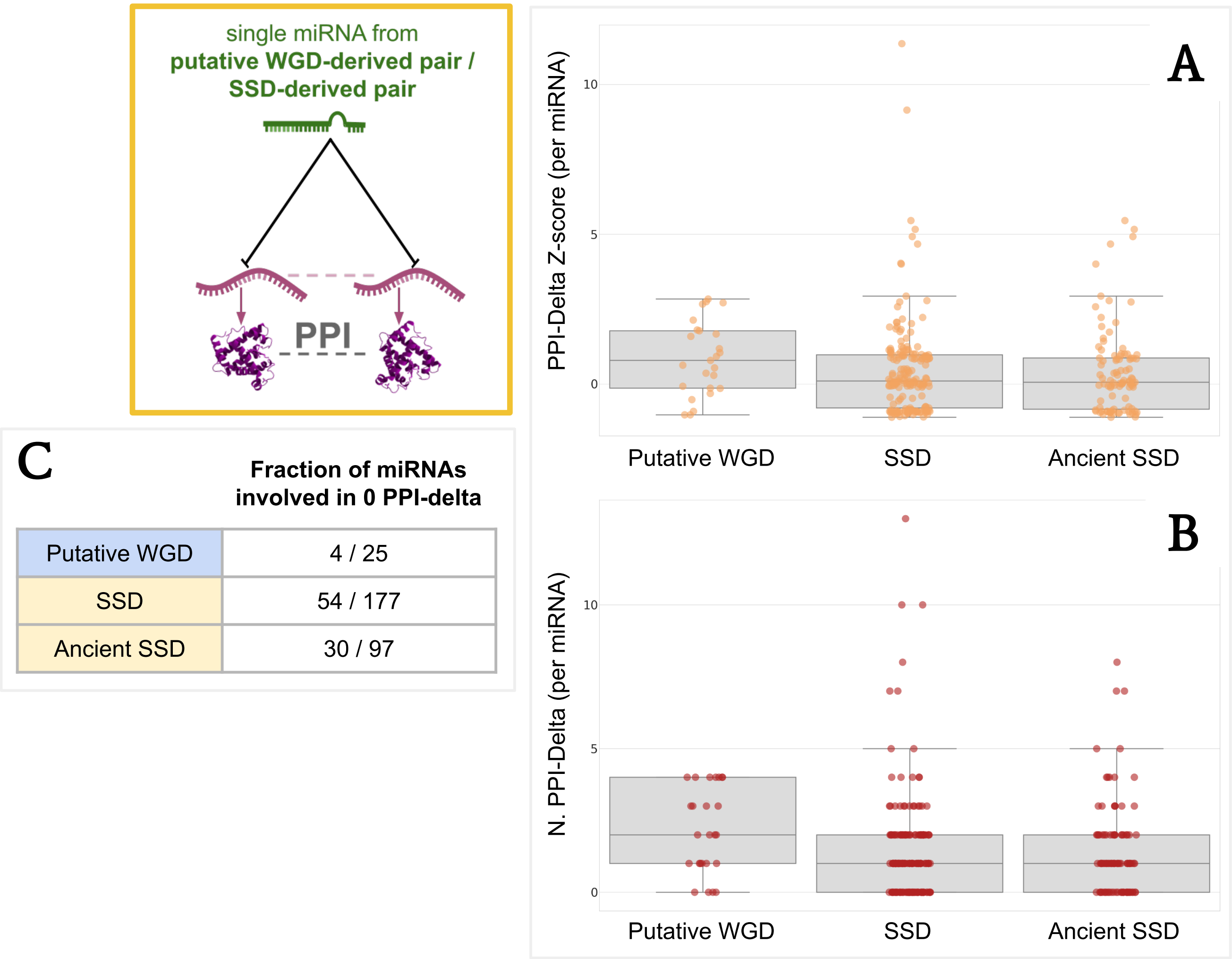


**Figure 10.** Same as 6 but changing the protein-protein interaction network to STRING. P-values from the comparison of the Z-score distributions: $P_{PutativeWGD\ vs\ SSD} = 0.068$, $P_{PutativeWGD\ vs\ Ancient\ SSD} = 0.059$ (Kolmogorov-Smirnov test).

#### 5 Discarded pairs from the Z-score plots

When plotting the Z-score for each duplicated miRNA pair, a few pairs are discarded because of the impossibility to compute a Z-score. This happens because, given a number of motif $n > 0$ observed in the real network, the average value $\overline{n}_{null}$ resulted in being equal to 0, and so also $\sigma_{null} = 0$. This rarely happens, but for the sake of completeness we report here the pairs discarded for each plot reported both in the Main Text (Section 1.3) and the Supplementary material (Sections 2 and Section 3) where this happened, together with the number of motif ($n$) they are involved in the real network.

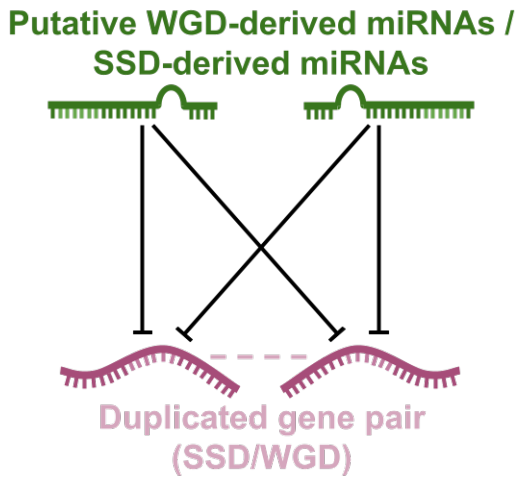


| TarBase | | |
|---|---|---|
| Pair | Set | $n$ |
| MIR135A1 - MIR135A2 | *SSD* and *Ancient SSD* | 1 |
| MIR302A - MIR302B | *SSD* | 1 |
| MIR302A - MIR302D | *SSD* | 1 |
| MIR518B - MIR518D | *SSD* | 1 |

**Table 5.** Discarded pairs, with respective number $n$ of motifs, in the analysis of simple bifan motif in the TarBase network (Section 1.3.3, Main Text).

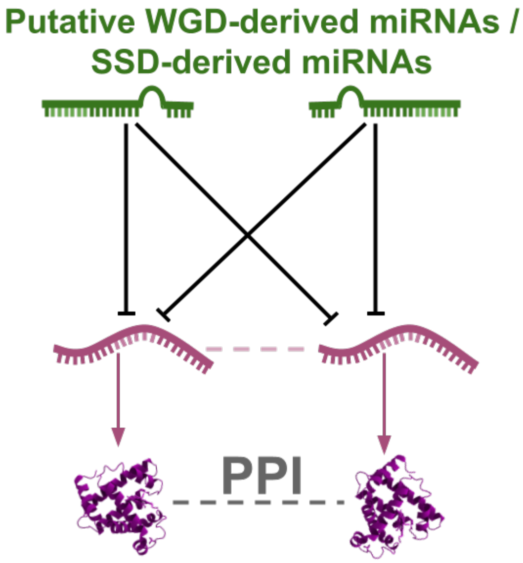


| **A.** TarBase + PrePPI | | |
|---|---|---|
| Pair | Set | *n* |
| MIR128-1 - MIR128-2 | *WGD* | 1 |
| MIR100 - MIR99A | *SSD* and *Ancient SSD* | 1 |
| MIR124-2 - MIR124-3 | *SSD* and *Ancient SSD* | 2 |

| **B.** TarBase + STRING | | |
|---|---|---|
| Pair | Set | *n* |
| MIR124-2 - MIR124-3 | *SSD* and *Ancient SSD* | 1 |

| **B.** MirDIP + STRING | | |
|---|---|---|
| Pair | Set | *n* |
| MIR320B1 - MIR320C2 | *SSD* | 1 |
| MIR518E - MIR526A1 | *SSD* | 1 |

**Table 6.**
**A.** Discarded pairs, with respective number *n* of motifs, in the analysis of PPI-bifan motif in the TarBase + PrePPI networks (Section 1.3.4, Main Text).
**B.** Discarded pairs, with respective number *n* of motifs, in the analysis of PPI-bifan motif in the MirDIP + STRING networks (Section 4 Fig.9, Supplementary).

### 6 Summary of the presence of duplicated genes and miRNAs TarBase and MirDIP

The presence of a miRNA/gene of interest in a given miRNA-Gene interaction network or a protein-protein interaction network is not granted, because many factors (biological, technical etc.) may lead to the absence of a certain number of elements. We provide here a summary of the presence of duplicated miRNAs and duplicated genes inside all the networks employed in the Main Text. Individual genes and individual miRNAs from different sets overlap (since they may have undergone subsequent duplication of different kinds, like the example in Section 2.5 of the Main Text)

#### 6.1 miRNA-target interaction networks: TarBase and MirDIP

| | Total | TarBase | MirDIP |
|---|---|---|---|
| putative intragenic Ohno-miRNAs (*WGD*) | 30 | 28 (93%) | 30 (100%) |
| Ensembl paralogue miRNAs (*SSD*) | 191 | 168 (88%) | 191 (100%) |
| Ensembl paralogue miRNAs pre-Sarcopterygii (*Ancient SSD*) | 111 | 108 (97%) | 111 (100%) |

**Table 7.** Number and percentage of single relevant miRNAs present in different interaction networks.

| | Total | TarBase | MirDIP |
|---|---|---|---|
| putative intragenic Ohno-miRNA pairs (*WGD*) | 20 | 18 (90%) | 20 (100%) |
| Ensembl paralogue miRNA pairs (*SSD*) | 685 | 411 (60%) | 685 (100%) |
| Ensembl paralogue miRNA pairs pre-Sarcopterygii (*Ancient SSD*) | 174 | 172 (99%) | 174 (100%) |

**Table 8.** Number and percentage of relevant miRNA pairs present in different interaction networks. A pair is counted as present only if both miRNAs are present in the interaction network.

| | Total | TarBase | MirDIP |
|---|---|---|---|
| Ohnolog genes (*WGD*) | 7,776 | 4,480 (58%) | 6,915 (89%) |
| SSD-derived genes (*SSD*) | 14,054 | 6,582 (48%) | 10,941 (79%) |
| SSD-derived genes pre-Sarcopterygii (*Ancient SSD*) | 13,162 | 6,421 (49%) | 10,523 (80%) |

**Table 9.** Number and percentage of single relevant protein-coding genes present in different interaction networks.

| | Total | TarBase | MirDIP |
|---|---|---|---|
| Ohnolog gene pairs (*WGD*) | 9,870 | 2,982 (32%) | 7,482 (80%) |
| SSD-derived gene pairs (*SSD*) | 132,003 | 29,008 (24%) | 71,713 (58%) |
| SSD-derived gene pairs pre-Sarcopterygii (*Ancient SSD*) | 121,043 | 28,621 (23%) | 69,973 (58%) |

**Table 10.** Number and percentage of relevant protein-coding gene pairs present in different interaction networks. A pair is counted as present in an interaction network only if both genes are present.

##### 6.2 Protein-protein interaction networks: PrePPI and STRING

| | Total | PrePPI | STRING |
|---|---|---|---|
| Ohnolog genes (*WGD*) | 7,776 | 4,364 (56%) | 4,151 (53%) |
| SSD-derived genes (*SSD*) | 14,054 | 6,481 (46%) | 6,729 (48%) |
| SSD-derived genes pre-Sarcopterygii (*Ancient SSD*) | 13,162 | 6,329 (48%) | 6,428 (49%) |

**Table 11.** Number and percentage of single relevant protein-coding genes present in different protein-protein interaction networks.

| | Total | PrePPI | STRING |
|---|---|---|---|
| Ohnolog gene pairs (*WGD*) | 9,870 | 3,491 (35%) | 3,166 (32%) |
| SSD-derived gene pairs (*SSD*) | 132,003 | 31,485 (24%) | 23,805 (18%) |
| SSD-derived gene pairs pre-Sarcopterygii (*Ancient SSD*) | 121,043 | 30,835 (25%) | 22,540 (19%) |

**Table 12.** Number and percentage of relevant protein-coding gene pairs present in different protein-protein interaction networks. A pair is counted as present in an interaction network only if both genes are present.

#### 7 Putative ohnolog miRNAs in the mouse genome

Here we report the lists of the putative intragenic ohnolog miRNAs in the mouse genome, whose sequence similarity is plotted in Fig.4 of the Main Text. Pairs highlighted in red are recognized as duplicated pairs by Ensembl.

**Table 13.** Putative ohnolog miRNA pairs with their respective host genes

| Putative Ohno-miRNA 1 | Host gene 1 | Putative Ohno-miRNA 2 | Host gene 2 |
|---|---|---|---|
| Mir103-1 | Pank3 | Mir103-2 | Pank2 |
| Mir26b | Ctdsp1 | Mir26a-1 | Ctdspl |
| Mir26b | Ctdsp1 | Mir26a-2 | Ctdsp2 |
| Mir26a-1 | Ctdspl | Mir26a-2 | Ctdsp2 |
| Mir199b | Dnm1 | Mir199a-1 | Dnm2 |
| Mir199b | Dnm1 | Mir199a-2 | Dnm3 |
| Mir1991-1 | Dnm2 | Mir199a-2 | Dnm3 |
| Mir218-1 | Slit2 | Mir218-1 | Slit3 |
| Mir152 | Copz2 | Mir148b | Copz1 |
| Mir103-1 | Pank3 | Mir107 | Pank2 |
| Mir103-2 | Pank2 | Mir107 | Pank1 |
| Mir211 | Trpm1 | Mir204 | Trpm3 |
| Mir128-2 | Arpp21 | Mir128-1 | R3hdm1 |
| Mir208a | Myh6 | Mir499 | Myh7b |
| Mir208b | Myh7 | Mir499 | Myh7b |

**Table 14.** SSD-derived paralogue miRNA pairs with their respective host genes.

| Putative paralogue miRNA 1 | Host gene 1 | Putative paralogue miRNA 2 | Host gene 2 |
|---|---|---|---|
| Mir208a | Myh6 | Mir208b | Myh7 |
| Mir452 | Gabre | Mir105 | Gabra3 |
| Mir582 | Pde4d | Mir139 | Pde2a |
| Mir873b | Lingo2 | Mir218-2 | Slit3 |
| Mir873b | Lingo2 | Mir218-1 | Slit2 |
| Mirlet7f-2 | Huwe1 | Mir140-1 | Wwp2 |
| Mir455 | Col27a1 | Mir676-1 | Eda |
| Mir3060 | Rnf215 | Mir340 | Rnf130 |
| Mir196a-2 | Hoxc5 | Mir10b | Hoxd4 |
| Mir1962 | Ptpre | Mir153 | Ptprn2 |

Some pairs are obviously missing, their absence is attributable to wrong annotations in the different genomes. To cite an example, let's consider the pair MIR10A-MIR10B in the human genome, recognized by our analysis as a putative ohnolog pair. Both MIR10A and MIR10B have orthologues in the mouse genome (Mir10a and Mir10b) located in the Hox cluster. However, Mir10a is not reported to be intragenic, it's instead located less than 10,000 bases upstream the Hoxb3 gene, thus preventing it from being recognized by our pipeline (conversely, Mir10b is correctly hosted on Hoxd3).
Some pairs are correctly missing, as the well-known MIR33A-MIR33B pair. While Mir33 in the mouse genome is correctly located within the Srebf2 gene, the Srebf1 lacks the orthologue of MIR33B. However, there are many vertebrates whose genomes conserve both the orthologues of MIR33A and MIR33B, for example the Green Anole (*Anolis Carolinensis*) genome preserve both miRNAs, despite being phylogenetically more distant from the human genome than the mouse.

#### 8 An updated compendium of *bona fide* ohnolog miRNA pairs

| miRNA 1 | miRNA 2 | Type | Source |
|---|---|---|---|
| MIR199A1 | MIR199B | Intragenic | * |
| MIR199A1 | MIR199A2 | Intragenic | * |
| MIR199B | MIR199A2 | Intragenic | * |
| MIR103A1 | MIR103A2 | Intragenic | * |
| MIR103A1 | MIR107 | Intragenic | * |
| MIR103A2 | MIR107 | Intragenic | * |
| MIR26B | MIR26A1 | Intragenic | * |
| MIR26B | MIR26A2 | Intragenic | * |
| MIR26A1 | MIR26A2 | Intragenic | * |
| MIR196A2 | MIR196A1 | Intragenic | * |
| MIR10A | MIR10B | Intragenic | * |
| MIR218-1 | MIR218-2 | Intragenic | * |
| MIR204 | MIR211 | Intragenic | * |
| MIR152 | MIR148B | Intragenic | * |
| MIR128-1 | MIR128-2 | Intragenic | * |
| MIR153-1 | MIR153-2 | Intragenic | * |
| MIR33B | MIR33A | Intragenic | *, [1] (relaxed) |
| MIRLET7A3 | MIRLET7E | Intergenic | [1] (strict) |
| MIRLET7A3 | MIRLET7F2 | Intergenic | [1] (strict) |
| MIRLET7F2 | MIRLET7A2 | Intergenic | [1] (strict) |
| MIRLET7A3 | MIRLET7A2 | Intergenic | [1] (strict) |
| MIR124-1 | MIR124-2 | Intergenic | [1] (strict) |
| MIRLET7A3 | MIRLET7A1 | Intergenic | [1] (intermediate) |
| MIRLET7E | MIRLET7A2 | Intergenic | [1] (intermediate) |
| MIRLET7A1 | MIRLET7A2 | Intergenic | [1] (intermediate) |
| MIR124-1 | MIR124-3 | Intergenic | [1] (intermediate) |
| MIRLET7C | MIRLET7A2 | Intergenic | [1] (intermediate) |
| MIR133A1 | MIR133A2 | Intergenic | [1] (intermediate) |
| MIR133A1 | MIR133B | Intergenic | [1] (intermediate) |
| MIRLET7F2 | MIRLET7A1 | Intergenic | [1] (intermediate) |
| MIR7-2 | MIR7-3 | Intergenic | [1] (intermediate) |
| MIR7-1 | MIR7-2 | Intergenic | [1] (intermediate) |
| MIR1-1 | MIR1-2 | Intergenic | [1] (intermediate) |
| MIR206 | MIR1-2 | Intergenic | [1] (intermediate) |

| miRNA 1 | miRNA 2 | Type | Source |
|---|---|---|---|
| MIRLET7A3 | MIRLET7C | Intergenic | [1] (relaxed) |
| MIR7-1 | MIR7-3 | Intergenic | [1] (relaxed) |
| MIRLET7E | MIRLET7C | Intergenic | [1] (relaxed) |
| MIR196A2 | MIR96B | Intergenic | [1] (relaxed) |
| MIR1-1 | MIR206 | Intergenic | [1] (relaxed) |
| MIRLET7C | MIRLET7F2 | Intergenic | [1] (relaxed) |
| MIRLET7C | MIRLET7A1 | Intergenic | [1] (relaxed) |
| MIR9-1 | MIR9-3 | Intergenic | [1] (relaxed) |
| MIR9-1 | MIR9-2 | Intergenic | [1] (relaxed) |
| MIRLET7E | MIRLET7A1 | Intergenic | [1] (relaxed) |
| MIR181A1 | MIR181A2 | Intergenic | [2] |
| MIR181B1 | MIR181B2 | Intergenic | [2] |

**Table 15.** List of *bona-fide* ohnolog miRNA pairs from different sources. The asterisk ("*") indicates that the miRNAs have been reported as putative ohnologs in this publication.
